# Structural and functional characterization of allatostatin receptor type-C of *Thaumetopoea pityocampa* revealed the importance of Q271^6.55^ residue in G protein-dependent activation pathway

**DOI:** 10.1101/2020.07.01.174037

**Authors:** Aida Shahraki, Ali Isbilir, Berna Dogan, Martin J. Lohse, Serdar Durdagi, Necla Birgul-Iyison

## Abstract

Insect neuropeptide receptors are among the potential targets for designing next-generation pesticides. Activation of allatostatin receptor type C (AstR-C), a G Protein-coupled receptor (GPCR), upon stimulation with its endogenous ligand, allatostatin C (AST-C), leads to the inhibition of juvenile hormone (JH) secretion that consequently regulates physiology of insects. Here we conducted *in silico* and *in vitro* approaches to characterize the structure and function of AstR-C of *Thaumetopoea pityocampa (T*.*pit)*, a well-known pest in Mediterranean countries. The sequence of AstR-C and AST-C were derived from whole genome sequencing (WGS) data. Resonance energy transfer (RET) methods were used to investigate the downstream effectors of the receptor and the temporal kinetics of G protein activation. Three-dimensional (3D) structure of AstR-C constructed via homology modeling methods was subjected to molecular dynamics (MD) simulations and docking studies to identify the orthosteric pocket. Our results showed that *T*.*pit* AstR-C couples to Gαi/o subtype of G proteins at sub-nanomolar ranges of the the ligand with the G protein recruitment and activation kinetics of ∼4 and 6 seconds, respectively, when 1 nM AST-C is administered. At the increasing concentration of native ligand, βarrestin was shown to be recruited at nanomolar ranges the ligand. Docking and MD simulation studies revealed the importance of extracellular loop 2 (ECL2) in *T*.*pit* AstRC/AST-C interaction, and combination of *in silico* and *in vitro* methods supported the accuracy of the built model and the predicted orthosteric pocket. Q271^6.55^ (Ballesteros-Weinstein generic numbering) was found to have a substantial role in G protein dependent activation of AstR-C possibly via contributing to the flexibility of the structure.

## 1. Introduction

Allatostatins (ASTs), are pleiotropic peptides abundant in arthropods. Allatostatin C (AST-C) is recognized by PISCF stretch of amino acids in its C-terminus. This subtype was first found in the brain of *Manduca sexta* that belongs to Lepidoptera family^1^. ASTs play a crucial role in modulating the physiology of insects and crustacean species due to their inhibitory effect on the synthesis of juvenile hormone (JH)^2^. Phenotypic traits, physiological and developmental processes of insects are dominated by this lipid-like hormone^3^. AST-C exert its downstream effect upon binding to the cognate receptor which belongs to G protein-coupled receptors (GPCRs). Thus, neuropeptide GPCRs are proposed as ideal targets for the development of novel anti-parasite agents and insecticides in veterinary medicine and agriculture^4^. GPCRs are seven-transmembrane-domain (TM) proteins present in different organisms from bacteria to fungi and animals responding to a plethora of extracellular stimulations^5^. Human GPCRs are under extensive investigation because of their importance in drug discovery studies. But insect GPCRs despite their significance in proposing new mode-of-action for the development of next generation pesticides have not been studied well and overlooked^6, 7^. Characterization of novel GPCRs in insects by investigating their structure, downstream effectors and binding pocket can provide valuable information that can be utilized for developing new molecules targeting neuropeptide receptors of insects and controlling the physiological processes of harmful insects.

*Thaumetopoea pityocampa* (*T.pit*) (Lepidoptera: Thaumetopoeidae), pine processionary moth, is one of the most serious pests residing in South Europe, North Africa and Mediterranean countries^8^. This insects feed from the leaves of pine trees and their outbreaks can cause severe defoliation of pine forests. Gregarious, urticating larvae are responsible for severe public and animal health concern as they can cause dermatitis and other severe allergic responses^9, 10^.

In this study, our aim was to characterize the structure and function of *T.pit* allatostatin receptor type C (AstR-C). To this goal, the sequence of AstR-C and its endogenous ligand, AST-C, were derived from whole genome sequencing (WGS) data of *T.pit* that is sequenced and analyzed in our lab for the first time. Förster-and bioluminescence resonance energy transfer (FRET and BRET) methods were used to investigate the downstream effectors of the receptor. The structures of AstR-C and AST-C were investigated using state-of-the-art *in silico* approaches. The orthosteric pocket of the receptor was identified combining molecular dynamics (MD) simulations and docking studies, and it was further validated by *in silico* and *in vitro* approaches. Q271 at 6.55 position (Ballesteros-Weinstein numbering)^11^ was identified to have profound effect on the activation of the receptor and MD simulations analysis revealed its importance on the flexibility of the structure.

## 2. Results and Discussion

### 2.1. *T.pit* AST-C

Sequence of *T.pit* AST-C was derived from the WGS data that is deposited in NCBI (*Thaumetopoea pityocampa* ASH_NBICSL_2017, Accession number of the assembled genome: WUAW00000000). The gene contained two introns that were removed and translated into the precursor protein. The obtained sequence was compared to AST-C preprohormone of other species (Supplementary Figure S1, panel A). The precursor neuropeptide needed to be further processed to obtain the mature peptide as neuropeptides in insects are produced as long precursors that process in the endoplasmic reticulum to make the bioactive peptide. In general, one precursor can results in different mature peptides^12^, but in case of AST-C, only one peptide produces from the precursor^2^. SignalP-5.0 was used to identify the signal peptide, and dibasic cleavage of the preprohormone was determined according to the rules provided by Veenstra^13^ (Supplementary Figure S1, panel B). The final neuropeptide was a 15 amino acids long peptide with sequence of QVRFRQCYFNPISCF. Glutamine in the N-terminus was converted to pyroglutamate based on the known post-translational modifications of AST-C ^14^. The sequence of the mature AST-C was compared with other lepidopteran and the sequences were found to be 100% identical ^1, 15, 16^.

AST-C is a relatively larger peptide that could fold into specific conformations and adopt secondary conformations. However, for many molecular modeling programs predicting the structure of such a large peptide from beginning *(ab initio*) could be challenging and most likely inconclusive. Hence, instead of trying to construct the 3D structure of Ast-C using modeling programs, homology modeling was performed using I-TASSER to obtain the structure of the ligand. Based on the known structural characteristics of the peptide, some particular considerations were considered in the modeling procedure. For instance, there is a structurally and functionally important disulfide bond between the 7^th^ and 14^th^ cysteine residues of AST-C^14^ and this bond was introduced in the final model by setting the distance restraint of 2.05 Å (i.e., the required distance for the formation of disulfide bond) between the two sulfur atoms. The constructed model showed a C-score of −1.16. C-score is the scoring system used by I-TASSER, and values higher than −1.5 are expected to possess the more probable folding of the protein^17^ so the constructed model here was in the acceptable range. The final structure of the ligand was modified in the N-terminus, converting glutamine to pyroglutamate (Supplementary Figure S2). Unfortunately, no structural data for AST-C is available in databases so we could not validate the accuracy of our model, but exerting many homology modeling runs with the already explained constrains all resulted in a turn-like secondary structure of AST-C.

### 2.2. *T.pit* AstR-C belongs to Class A GPCRs

Sequence of the receptor was derived from the WGS data of *T.pit* as well. Investigating the sequence of the receptor in pfam online tool^18^, it was found that it belongs to seven transmembrane rhodopsin family GPCRs (Supplementary Figure S3, panel A). AstR-C possesses all the conserved residues and motifs available in class A GPCRs (Supplementary Figure S3, panel B). The only exception is position 6.30 (Ballesteros-Weinstein generic numbering) at which Glu residue is substituted with a His residue. E6.30 plays an important role in the activation of class A GPCRs as it forms an ionic lock with R3.50 and T6.34. The same exception is observed for opioid receptors as well, but histidine substitution in these receptors is shown not to affect the activation of these receptors since the hydrogen bond network between this residue and R3.50 and T6.34 is still present^20^.

Subcellular localization of the receptor was determined. *T.pit* AstR-C was cloned in SYFP plasmid to fuse the C-terminus of the receptor with yellow fluorescent protein (YFP). AstRC-SYFP construct was transfected in HEK-TSA cells and plasma membrane and nucleus were stained. Live cell confocal microscopy imaging showed that the fluorescence signal from YFP was predominantly co-localized with the cell membrane marker, suggesting that *T.pit* AstR-C mainly localized in the plasma membrane (Figure 1).

**Figure 1.**
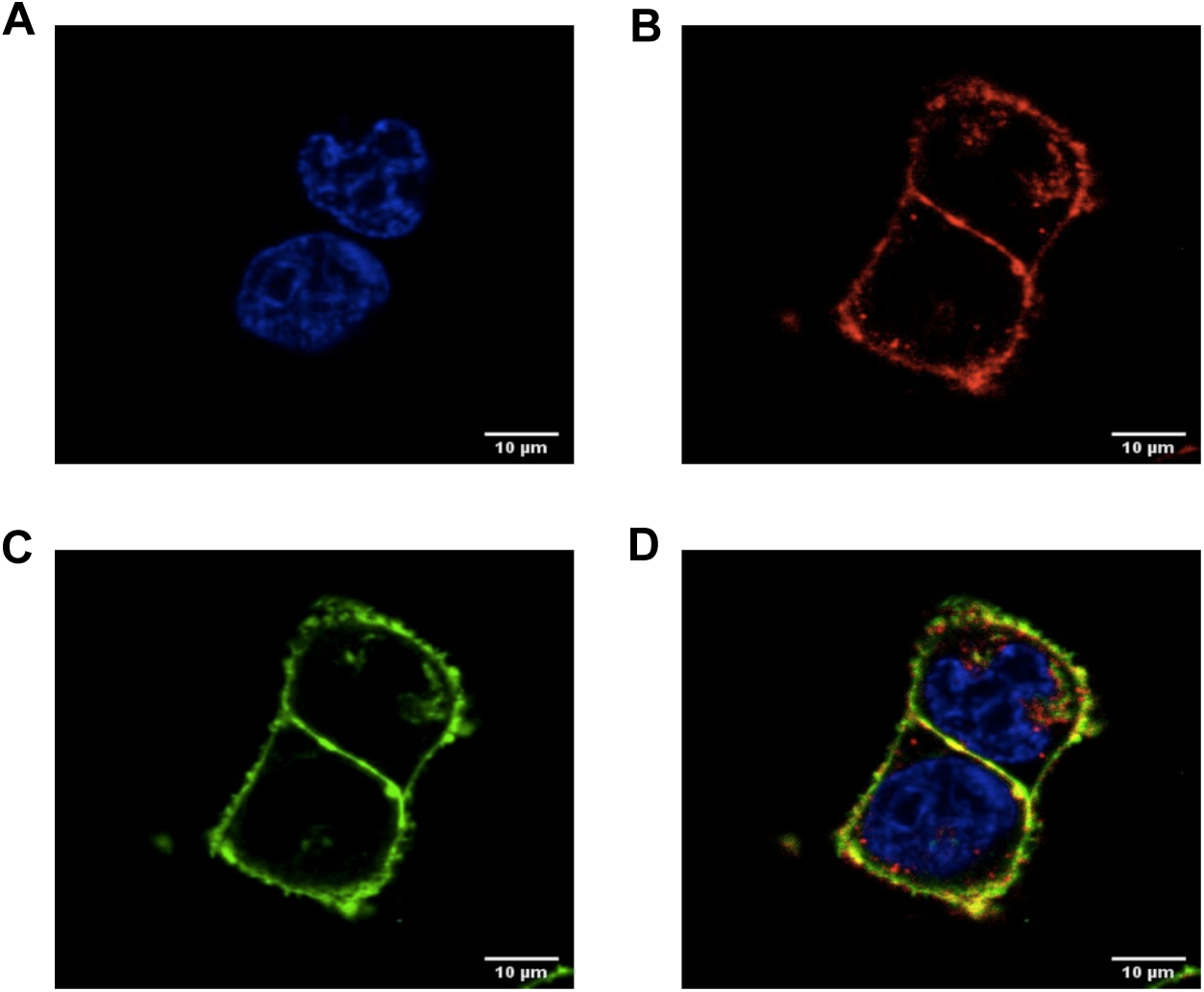
Cell localization of *T.pit* AstR-C. Confocal microscopy images of HEK-TSA cells transfected with AstRC-SYFP. (A) Nuclei is stained with Hoechst 33258 (blue). (B) Plasma membrane is stained with CellMask™ Deep red (red). (C) *T.pit* AstR-C (green). (D) Merged image obtained from the overlay of three images.

### 2.3. Downstream Effectors of *T.pit* AstR-C

GPCRs that are stimulated with their relevant ligand, in turn, could activate the intracellular G protein heterotrimers^5^. Four subtypes of these proteins are found in the cell, G_s_, G_i/o_, G_q/12_ and G_12/13_, and each initiate a specific downstream cascade^21^. To understand whether AST-C peptide can activate *T.pit* AstR-C, G protein activation assay was performed. Different biosensors were used to find the G protein subtype that couples to the receptor. G protein FRET biosensors were all tagged with YFP and cyan fluorescent protein (CFP) at γ- and α-subunits, respectively (Figure 2, panel A). FRET changes before and after application of the native ligand were measured. A decrease in FRET signal was expected provided that the receptor couples to the relevant G protein following the administration of the native ligand. The reduction happens since the distance between the donor (CFP) and acceptor (YFP) increases. In case of *T.pit* AstR-C, we observed a decrease in FRET signal when G_i2_ sensors were used. Hence, it was deduced that this insect neuropeptide receptor favors to be coupled to the G_i_ subtype (Figure 2, panel B). Three different G_i_ sensors were tested in this assay. Results showed that all G_i_ sensors (G_i1_, G_i2_ and G_i3_) couple to AstR-C with an EC_50_ at sub-nanomolar range but the highest ΔFRET shift was observed for G_i2_ (Figure 2, panel C). Therefore, in the following experiments G_i2_ was used.

**Figure 2.**
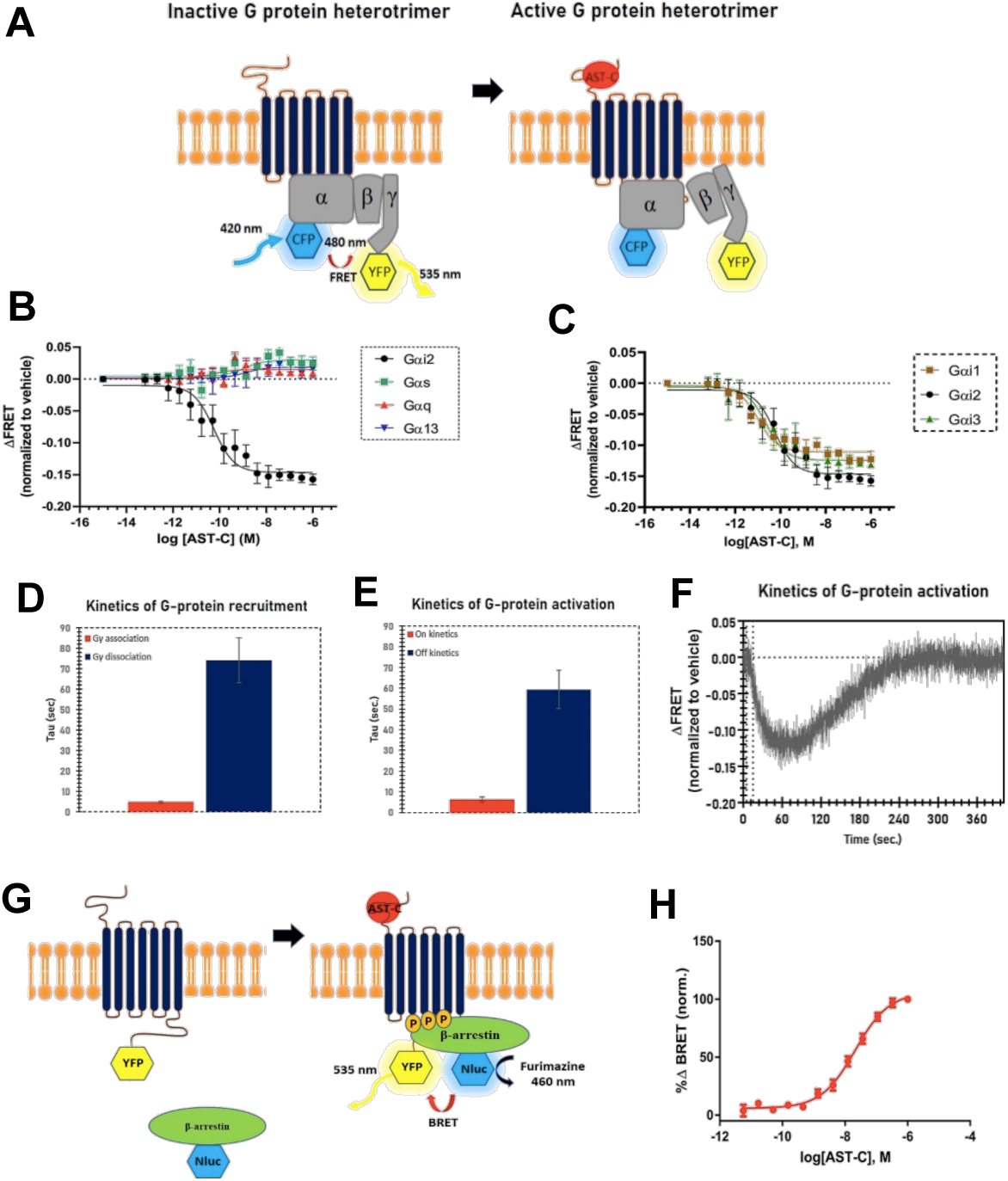
Downstream effectors of *T.pit* AstR-C. (A) Schematic representation of G-protein activation assay. (B) G protein activation of *T.pit* AstR-C in response to increasing concentrations of AST-C when different biosensors are transiently transfected. (C) G_i_ activation of the receptor in response to increasing concentrations of AST-C. (D) Kinetics of G-protein recruitment when *T.pit* AstR-C-SYFP and G_i2_ sensor are transiently transfected in the presence of 1 nM AST-C ligand. (E) Kinetics of G-protein activation when *T.pit* AstR-C-WT and G_i2_ sensor are transiently transfected in the presence of 1 nM AST-C ligand. (F) A representative trace of FRET response from a single HEK-TSA cell. (G) Schematic representation of βarrestin recruitment assay. (H) βarrestin recruitment to T.pit AstR-C in response to increasing concentrations of AST-C. Results from each 96-well plate experiment were normalized to max-min values from the same plate. Data was fit to Hill equation, using the four-parameter dose-response fit function of GraphPad Prism6. The presented data is representative for at least three different transfections performed on three experimental days. The error bars represent standard deviation (SD). Values of the bar graphs in the kinetics measurements are the average of data obtained from four cells and at least 3 independent experiment days. Values of the bar graphs in the dose response curves are the average of data obtained from at least three independently conducted experiments.

Kinetics studies were conducted at two events, G protein recruitment, and G protein activation. Kinetics of the recruitment of G protein complex to AstR-C was investigated to evaluate how fast the G protein complex is recruited to the receptor following a brief application of the ligand (10 second), and how long it takes to be dissociated from the receptor after washing it off (400 seconds). To this aim, the C-terminus of the receptor and the gamma subunit of G protein complex were tagged with YFP and CFP, respectively. In temporal kinetics of G protein recruitment experiment the off-kinetics was best measurable at 1 nM concentration of the native peptide, AST-C, during the total 400-seconds measurement time. On average, a τ-value of 4.7 and 74.1 second were yielded for the association and dissociation of G protein to the receptor at 1 nM concentration of AST-C, respectively (Figure 2, panel D).

The kinetics of G protein activation were investigated as well to evaluate the G protein activation kinetics. Fluorescent tags used in this experiment were identical to the ones used in G protein activation assay (Figure2, panel A). The experiment here shows the time that Gi protein remains active following a brief application of the ligand (*on* kinetics, G protein activation (Gα-Gβγ subunit rearrangement/dissociation)) and the time required for the G protein to return to its basal level (*off* kinetics, G protein deactivation (subunit rearrangement/reassociation)) when the ligand is being washed off by perfusing buffer (instead of ligand) to the cell. Complete inactivation of G protein was observable at 1 nm concentrations with the τ-value of 6.2 second for *on* kinetics of and 59.3 second for off kinetics (Figure 2, panel D and Figure 2, panel E).

Non-visual arrestins, βarrestin1 (arrestin-2) and βarrestin2 (arrestin-3), are cytosolic proteins that bind agonist stimulated receptors^22^. In this study, βarrestin2 recruitment to *T.pit* AstR-C upon the simulation with different concentrations of ligand was investigated using βarrestin with a C-terminally incorporated nanoluciferase NanoLuc ^23^ luciferase (19 kDa; Nluc) as the donor and YFP at the C-terminus of the receptor as the acceptor. Nluc emission happens in the presence of its substrate, furimazine (Figure 2, panel G). This assay showed that the βarrestin is recruited to the receptor at nanomolar range (EC_50_ values = 37 nM) (Figure 2, panel H).

### 2.4. 3D Structure of *T.pit* AstR-C

As there is no crystal structure of AstR-C of *T.pit*, 3D homology model of the receptor was built using SWISS-MODEL webserver (https://swissmodel.expasy.org). Different templates were used to build a reliable model, and the one constructed based on Mus-musculus Mu opioid receptor (PDB ID, 6DDE) with the resolution of 3.5 Å^24^ was chosen as it was resolved along with an agonist and human nucleotide-free Gi and more importantly, because the receptor was in the active state. This active template showed 37.15% sequence identity to *T.pit* AstR-C. The constructed model was subjected to short MD simulations (25-ns) to relax and refine the structure, and the stability of the built model was evaluated by investigating root mean square deviations (RMSD) and root mean square fluctuations (RMSF) observed during the MD simulation time (Supplementary Figure S4, panels A and B). To account for the possible role of the N-terminus of the receptor in ligand binding and orthosteric pocket formation, this part was modeled separately using I-TASSER webserver and merged with the model (Supplementary Figure S4, panel C and D). In addition, the quality of the model was evaluated by inspecting the Ramachandran’s plot (Supplementary Figure S4, panels E and F). In order to have a better prediction regarding the binding pocket of the receptor and obtaining the most active-like conformation of binding pocket, the final model was built as a complex of G_α_ and receptor in which α-subunit conformation of G protein heterotrimer was taken from the structure resolved in Mus-musculus μ-opioid receptor – G_α_-protein complex. There were two gap regions in the resolved G_α_ structure which were modeled using the Crosslink proteins module of Schrödinger based on the UniProt sequence (UniprotKB: P63096) (Supplementary Figure S5). The final system having *T.pit* AstR-C and G_α_ in the intracellular interface was subjected to three individual replicas of 500-ns MD simulations runs initiating with different velocity distributions. The stability of the system was evaluated using RMSD and RMSF plots. RMSD and RMSF analysis showed relatively high values that mainly stem from 36 N-terminus residues of AstR-C. This region localizes in the extracellular matrix and is highly flexible that consequently results in the increased values of RMSD and RMSF, but even if it was considered in the calculations, the RMSD reached to a plateau during the MD simulation time (Supplementary Figure S6, panels A, B, C and D). Besides the N-terminus, other parts of the receptor showed a good stability with low RMSD and RMSF fluctuations throughout the simulations. Residues with high fluctuation values are all residing in the extracellular and intracellular loops, ECL and ICL, respectively, and higher fluctuations for these loop regions are expected. A part of the C-terminus that was modeled in the final model also showed high fluctuations. The structure of G_α_ was also stable during the MD simulation, though being a cytosolic protein, it showed higher RMSD values in comparison to AstR-C (Supplementary Figure S6, panels E and F).

### 2.5. Orthosteric binding pocket of *T.pit* AstR-C

Molecular docking and MD simulations studies were combined to identify the orthosteric binding pocket of the receptor. Keeping the ligand, AST-C, flexible and the receptor rigid, protein-protein docking was performed in ClusPro webserver (https://cluspro.org). 1000 rotamers of AST-C were generated by the program and 946 of them clustered together in an identical pose. The best pose with the lowest energy of −1621.8 kcal mol^-1^ was chosen. This pose was subjected to 500-ns MD simulations. The Molecular Mechanics/Generalized Born Surface Area (MM/GBSA) analysis was conducted for 100 frames selected from the MD simulation trajectories to calculate the binding free energy of the ligand (ΔG). The average ΔG was calculated as −147.92 ± −15.88 kcal mol^-1^. The RMSD and RMSF deviations of the receptor and the ligand throughout the MD simulation time were evaluated as well. Two different RMSD fitting modes were considered to assess the stability of ligand AST-C. While the first one was the RMSD with respect to the first frame of the backbone atoms of the receptor to evaluate the translational motion of the ligand at the binding pocket, in the second fitting mode rotational motion or in other words internal fluctuations of the ligand at binding pocket was evaluated. We denoted the translational motion of ligand by “Lig-fit-Protein” RMSD and internal fluctuations by “Lig-fit-Ligand” RMSD. During the first 100-ns of the simulation time, the ligand showed high deviations from the first frame and then in the remaining time the fluctuations decreased and AST-C reached to a stable mode which continued until the end of the simulation time (Supplementary Figure S7, panel A). RMSF values of AstR-C showed higher fluctuation at loops, N-terminus and C-terminus, which was expected. Not considering the N-terminus, the highest fluctuation was observed for the ECL3 (Supplementary Figure S7, panels B and C). Except for the first 4 residues of the ligand, the ligand showed not much fluctuations and G_α_ RMSF values were high at the loop regions (Supplementary Figure S7, panel D and E). MD simulations trajectories were analyzed and the interaction between the ligand and the receptor was investigated to find the residues of AstR-C that mainly contribute to the formation of the orthosteric pocket as well as to identify significant residues in receptor-ligand interactions (Figure 3, panel A and B). Results showed that ECL2 takes the main role in the establishment of the interaction, and residues in this region forms one or more type of interaction with the ligand during the MD simulations time. ECL3 was also involved in the binding site. Hydrogen bonding interactions was found to be the prevalent type of interaction in the binding pocket, but hydrophobic interactions, salt-bridge and water bridge interactions were also involved (Figure 3, panel C and D).

**Figure 3.**
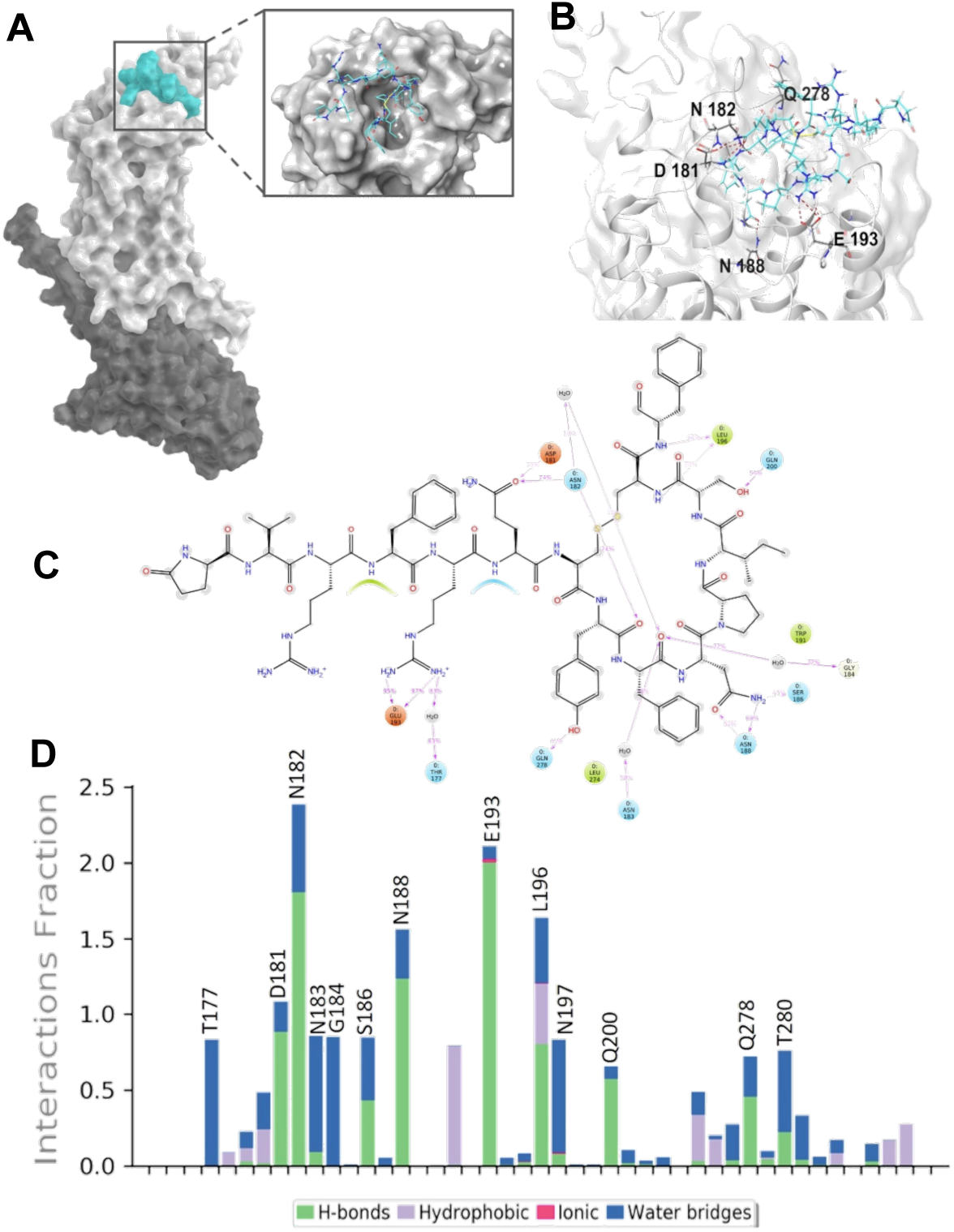
MD Simulation interaction analysis. (A) Surface representation of AST-C (colored Turquoise) at the binding pocket of AstR-C, represented in light gray. G□ is depicted in dark gray. (B) 3D ligand interactions diagram of AST-C at the binding site of AstR-C. (C) 2D representation of Protein-Ligand interaction. Residues of the receptor surrounding the ligand are represented with different colors each showing the type of the interaction. (D) Protein-Ligand interaction fraction diagram. Stacked bars show the type of the interactions each residue of the receptor makes with ligand during the MD simulation time. Residues are with interaction fraction value higher than 0.5 are specified.

To verify the binding pocket suggested by docking and MD simulations, some of the residues of the receptor that form long-lived interactions in MD simulations with the peptide Ast-C were mutated to Alanine (Ala) *in silico*. Additionally, a point mutation of Q271A was generated, due to the well-known importance of 6.55 position in ligand binding in other GPCR Class A receptors^25, 26, 27^. 200-ns MD simulations were run for mutant receptors at apo-form, and new docking poses were generated for all the mutants using ClusPro server. These holo forms were then subjected to 200-ns MD simulations with and without G_α_ subunit. To investigate the influence of point mutations on the state of the structure i.e. being active, inactive, or intermediate state, Δ*d* was calculated according to *gpcrdb* recommended measurement in which the distance between two pairs of residues are measured and then subtracted (http://docs.gpcrdb.org/structures.html) (Table 1). In class A GPCRs, Δ*d* below 2.0 Å shows a structure at inactive state, between 2 to 7.15 Å is related to intermediate states of the structure and values higher than 7.15 Å are attributed to structures at active state. At the apo form, wildtype (WT) receptor was found to be in an intermediate state during the MD simulations time, however, Ala-substitution at D181 and N182 positions moved the state toward inactive states in apo form. The mutation on residues E193A and Q278A lead completely the opposite behavior and resulted in structures at active state. The receptor remained at an intermediate state for N188A, Q200A and Q271A mutants. Binding of AST-C to the receptor increased the Δ*d* values in general, and expectedly, shifted the structures toward more active states (Table 1). This was significant for WT receptor, in particular, for which binding of the ligand transitioned the state from intermediate to active. In contrast to the general trend observed for WT and other mutant receptors, point mutation Q271A shifted the state of the receptor to inactive state. At holo form, it was shown that mutations introduced in the binding pocket change the ligand binding pose when compared to the WT receptor (Figure 4, panels A). While ATS-C was mainly positioned between ECL2 and ECL3 at WT receptor, it seemed that at mutant receptors the ligand moved more toward the funnel of the receptor. It can be explained in part by the reduced steric clash in the binding pocket following the Ala substitution, due to the smaller side chian of Ala compared to the substituted ones, that allow the ligand to move deeper in the receptor ortosteric cavity.

**Table 1.** Δd values and state of structure for WT and mutant receptors at Apo and Holo forms (with/without Gα).

| **Apo**<br>(Å) | WT | D181A | N182A | N188A | E193A | Q200A | Q271A | Q278A |
| --- | --- | --- | --- | --- | --- | --- | --- | --- |
| $d_2$ | $21.7 \pm 0.9$ | $19.3 \pm 0.6$ | $19.5 \pm 0.7$ | $22.5 \pm 0.8$ | $22.7 \pm 0.5$ | $18.7 \pm 1.2$ | $21.9 \pm 0.7$ | $23.2 \pm 0.5$ |
| $d_1$ | $18.3 \pm 0.9$ | $20.4 \pm 0.7$ | $21.0 \pm 0.9$ | $15.4 \pm 0.5$ | $14.5 \pm 0.5$ | $15.6 \pm 0.6$ | $15.3 \pm 0.5$ | $14.2 \pm 0.5$ |
| $\Delta d$ | 3.4 | -1.04 | -1.44 | 7.09 | 8.2 | 3.08 | 6.6 | 9.38 |
| State | Inter-mediate | Inactive | Inactive | Inter-mediate | Active | Inter-mediate | Inter-mediate | Active |
| **Holo**<br>(Å) | WT | D181A | N182A | N188A | E193A | Q200A | Q271A | Q278A |
| $d_2$ | $21.5 \pm 0.9$ | $24.6 \pm 1.1$ | $24.5 \pm 1.5$ | $25.5 \pm 1.5$ | $22.8 \pm 0.9$ | $27.6 \pm 1.4$ | $21.3 \pm 0.6$ | $22.6 \pm 0.8$ |
| $d_1$ | $14.6 \pm 0.6$ | $14.1 \pm 0.4$ | $14.5 \pm 0.9$ | $14.6 \pm 0.9$ | $14.3 \pm 0.5$ | $18.2 \pm 1.0$ | $19.5 \pm 0.7$ | $17.6 \pm 0.6$ |
| $\Delta d$ | 6.9 | 10.15 | 5.0 | 10.8 | 8.4 | 9.4 | 1.8 | 4.9 |
| State | Inter-mediate | Active | Inter-mediate | Active | Active | Active | Inactive | Inter-mediate |
| **Holo ($G\alpha$)**<br>(Å) | WT | D181A | N182A | N188A | E193A | Q200A | Q271A | Q278A |
| $d_2$ | $23.5 \pm 0.9$ | $21.4 \pm 0.7$ | $22.4 \pm 0.5$ | $25.1 \pm 1.0$ | $22.1 \pm 0.4$ | $22.7 \pm 0.8$ | $21.5 \pm 0.5$ | $21.5 \pm 0.6$ |
| $d_1$ | $12.0 \pm 0.5$ | $15.5 \pm 0.5$ | $14.2 \pm 0.3$ | $14.9 \pm 0.5$ | $13.7 \pm 0.3$ | $19.1 \pm 1.0$ | $18.2 \pm 0.6$ | $17.5 \pm 0.6$ |
| Δd | 11.5 | 5.8 | 8.2 | 10.5 | 8.4 | 3.6 | 3.3 | 4.0 |
| State | Active | Inter-mediate | Active | Active | Active | Inter-mediate | Inter-mediate | Inter-mediate |

**Figure 4.**
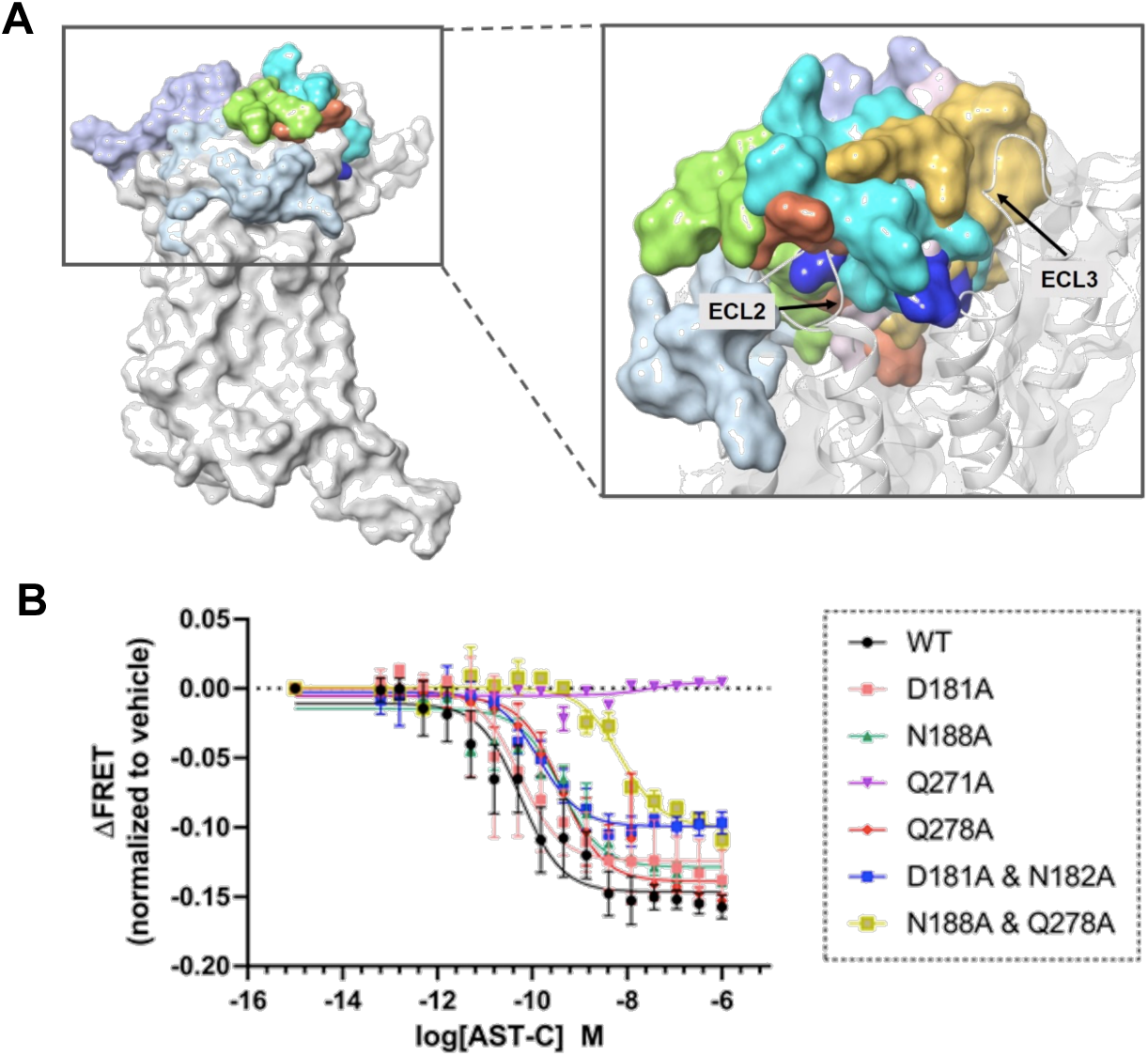
*In silico* and *in vitro* verification of the orthosteric pocket. (A) Effect of point mutations on the binding pose of AST-C was shown by superimposing receptors (WT and mutants) in holo form system. AST-C in WT is shown in turqoise. (B) Effect of mutations on dose-dependent G protein activation of *T.pit* AstR-C. The changes in FRET signal of mutant AstR-C were measured and compared with WT AstR-C upon the application of different doses of AST-C ligand. The data was fit to Hill equation, using the four-parameter dose-response fit function of GraphPad Prism6. The presented data is representative for at least three different transfections performed on three experimental days.

Following *in silico* studies, *in vitro* experiments were designed to validate the importance and significance of binding pocket-residing residues. The mutant receptors along with the WT receptor were tested in FRET-based G protein activation assay. Besides the point mutations, a combination of some of these point mutations were generated as well, in order to investigate the collective effect of these residues in forming stable contact with the ligand. In general, when receptor was mutated at the binding site, the dose-response curve shifted toward the higher concentration and higher EC_50_ values compared to that of WT AstR-C (Figure 4, panel B and Table 2). The observed effect implied the importance of these binding pocket residues in the activation of G proteins after coupling to the receptor. As expected, having more than one mutation in the binding pocket of the receptor led to more pronounced effects in the increase of EC_50_ values. In addition, the maximum response of AstR-C for G protein activation was decreased by almost 30% in double mutant receptors comparing to WT AstR-C further illustrating the significance of these residues in forming the orthosteric pocket of AstR-C and ligand binding interactions. No G protein activation was observed for Q271A mutant receptor. Overall, the results acquired here very well supported the accuracy of the built model and the predicted binding pocket.

**Table 2.** EC_50_ and R^2^ values of WT and mutant AstR-C compared to WT.

| Receptor Constructs | EC<sub>50</sub> (nM) | R<sup>2</sup> (Goodness of fit) |
| --- | --- | --- |
| WT | 0.057 | 0.89 |
| D181A | 0.053 | 0.77 |
| N188A | 0.39 | 0.95 |
| Q271A | - | 0.25 |
| Q278A | 0.40 | 0.91 |
| D181A & N182A | 0.13 | 0.93 |
| N188A & Q278A | 7.20 | 0.94 |

### 2.6. Q271A substitution

No direct interaction between Q271 residue and AST-C was observed in MD simulations of AstR-C and AST-C. However, *in silico* and *in vitro* analysis performed for the verification of the identified binding pocket, showed the drastic effect of Q271A substitution on receptor activation. Thus, we decided to investigate further the structural implication of this point mutation. First, we confirmed the membrane localization of this mutant (Supplementary Figure S8). Eliminating the possibility of not being localized in plasma membrane, we speculate that there are two possible scenarios for the observed effect of Q271A on the activation of the receptor. First, the point mutation might change the structure at apo form so that the ligand does not bind to the structure, and the second possibility is that the ligand binds to the receptor, but the structure cannot go through the conformational changes required for the G protein coupling and following activation. We tested both possibilities. MD simulations were performed both in Apo and Holo forms, and it was checked to see if a new binding pose could be obtained by CLusPro docking of mutated Apo form Q271A.

Superimposition of WT receptor and mutants at apo form revealed a distinct conformation of TM6 and ICL3 at mutant Q271A receptor, with more inwardly positioned TM6 and ICL3, suggesting a possible role of this point mutation on the overall structure of the receptor halting the conformational changes required for the activation. It is of note that all other receptor constructs including WT exploited similar conformations in this region when compared to each other (Supplementary Figure S9). Internal movements and displacements of the structures were more scrutinized by performing principle component analysis (PCA) and dynamical cross-correlation analysis to the trajectories of 500-ns MD simulations runs performed for WT and Q271A at apo and holo forms. First three principal components (PCs) covered more than 60% of all the movements in all systems (Supplementary Figure S10). Investigating the fluctuations of the structures in the first three PCs, it is obvious that Q271A mutation reduces the internal movements, especially in the ICL3 (Supplementary Figure S11). Comparing the eigenvalue magnitudes of PCs between WT and Q271A receptor, higher level of fluctuations was observed in all forms of WT, and eigenvalue magnitudes of Q271A were significantly lower (Figure 5). GPCRs are highly flexible allosteric proteins that their activation requires many conformational changes in the structure following the ligand binding but as PC analysis revealed, Q271A mutant receptor has considerably less internal motions that we speculated might be the underlying factor for the loss of activation. The trajectories were also investigated with cross-correlation analysis. The dynamical cross-correlation map (DCCM) showed that binding of the ligand to WT receptor results in the decrease in the population of un-correlated motions, shown in blue (Figure 6). This is more obvious for residues of ECL2, especially between 180 to 200 residues, in which our results showed their importance in protein-ligand interactions. However, this trend was not observed for Q271A mutant receptor. In fact, for this point mutation, the structure showed very different pattern of movement with si G_α_, the pattern was drastically different from the WT receptor and especially in ECL2 no correlated movement was detected. PCA and cross-correlation analysis together showed the internal motion changes that happens in the structure of the receptor following Ala substitution at Q271^6.55^.

**Figure 5.**
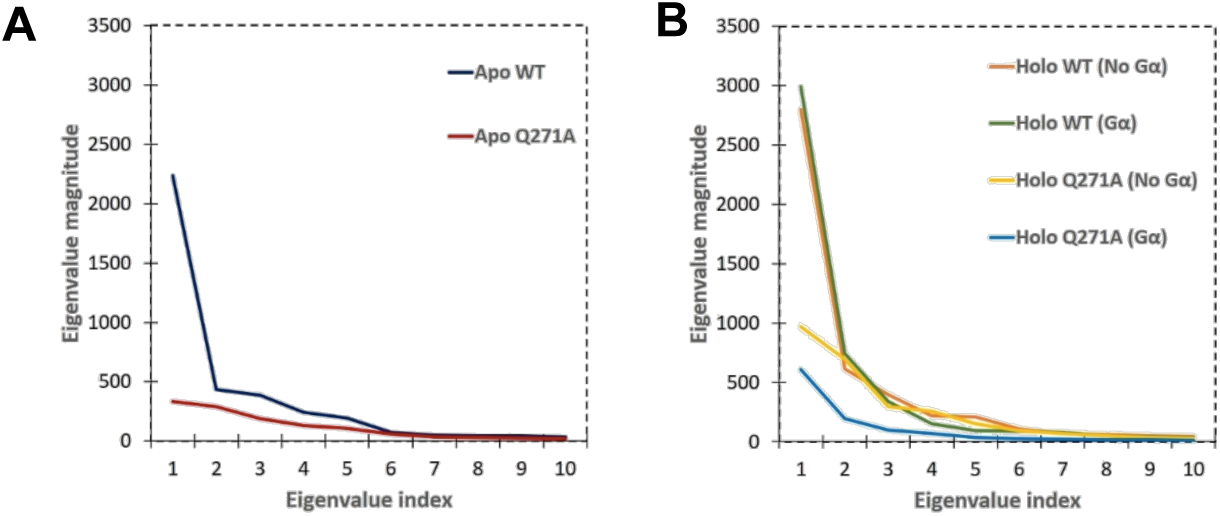
Eigenvalue magnitudes. Analysis of Eigenvalues corresponding to eignevalue indexes for of the first 10 modes of action of (A) WT and, (B) Q271A receptors at different states.

**Figure 6.**
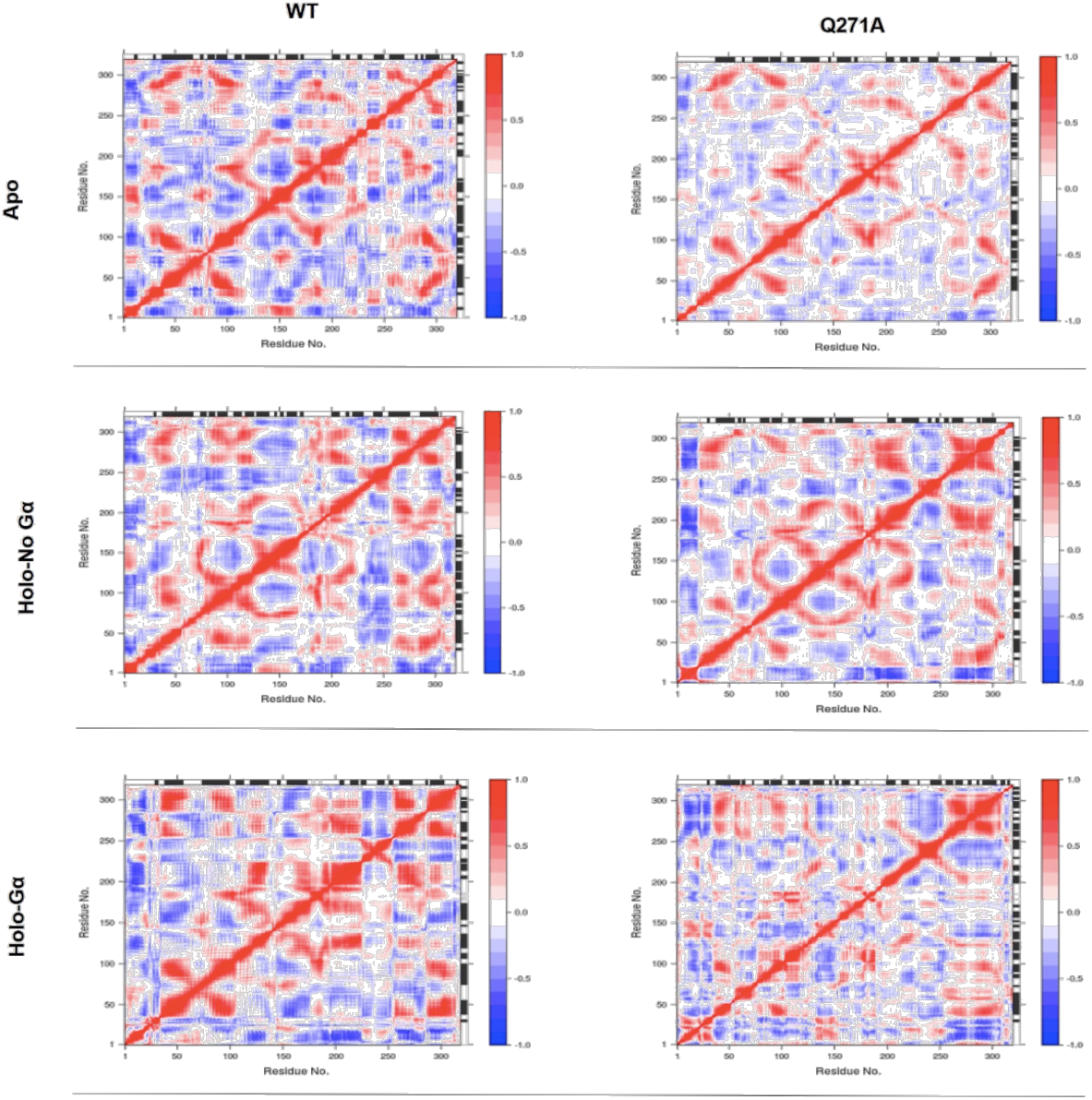
Dynamical cross-correlation map. Correlated (red) and un-correlated (blue) displacements were compared between WT and Q271A receptor at different states.

## 3. Conclusions

Upon binding to their cognate receptors, ASTs inhibit the secretion of JH that, in turn, regulates the downstream physiology in insect. Here, we conducted extensive *in silico* and *in vitro* studies to characterize the structure and function of AstR-C of pine processionary moth, a predominant pest residing in Mediterranean countries, South Europe, and North Africa. We derived the sequence of the receptor and endogenous ligand, AstR-C and AST-C, from the WGS data sequenced and analyzed in our lab. G protein recruitment and activation was observed via *T.pit* AstRC activation by sub-nanomolar concentrations of AST-C. This was anticipated as AstR-C in insects are ortholog of somatostatin receptors that signal via Gi/o G-protein subtype^28^. βarrestin, as another downstream effector was shown to be recruited to the receptor at nanomolar ranges. We investigated the temporal kinetics of G protein dependent signaling in the recruitment and activation steps, and our results showed that at 1 nM concentration of AST-C, it takes 4.7 seconds for G protein complex to be recruited to the receptor, and 6.2 seconds be dissociated to G_α_ and G_βγ_. Compared to other GPCRs activated by small molecules, the acquired activation time is longer which can be attributed to the large size of the peptide ligand, AST-C, and its binding modes ^29, 30, 31, 32^. Structural studies were resulted in a reliable 3D model of *T.pit* AstR-C and AST-C generated via homology modeling approaches. We then combined classic MD simulations and docking studies to predict the orthosteric pocket of the receptor. Investigating the trajectories obtained from multiple independent MD simulation runs, we identified residues of the receptor with main contribution in receptor-ligand interaction. In line with the literature of class A GPCRs, our results revealed the essential role of ECL2 in forming the binding cavity and ligand binding ^33^. ECL3 was also involved to lower extent. The significance of some residues selected from ECL2 and ECL3 in the formation of orthosteric pocket and activation of *T.pit* AstR-C was validated by *in silico* and *in vitro* methods. As a result of these studies we identified a residue at position 6.55 (Q271), which has no direct interaction with the ligand but has a critical role in AstRC activation. Ala substitution at Q271^6.55^ was found to be detrimental for the G protein activation pathway. Looking at atomic level, we showed that this mutation disrupts the internal movements of the receptor and changes the pattern of the correlated and un-correlated motions of residues when compared to the WT receptor. We attributed the significantly lower eigenvalues magnitudes of Q271A mutant at apo and holo forms to the lower level of flexibility in this mutated form, which in turn, blocks the conformational changes required for the GPCR activation. It is of note that a similar effect at the same position is reported by Change *et al*. ^34^, in kappa opioid receptor (κOR), where an Ala substitution disrupts the TM6 and ICL3 outward movements. Taken together, we believe that the characterization studies performed on the novel insect neuropeptide receptor, AstR-C of *T.pit*, and the structural and functional insights obtained here will positively contribute to the future studies aiming to exploit the potential of insect neuropeptide receptors in designing more environmentally friendly pest control agents.

## 4. Materials and Methods

### 4.1. AstR-C and AST-C sequences

The nucleotide sequence of AstR-C and AST-C were derived from the whole genome sequencing data of the insect performed by our group (*Thaumetopoea pityocampa* ASH_NBICSL_2017, accession number of the assembled genome: WUAW00000000). Protein sequences of AstR-C and AST-C of *Drosophila melanogaster* and *Helicoverpa armigera* were used as queries to search for their orthologs in the new assembly contigs of *T.pit*. Using these queries, NCBI-tblastn was performed at default parameters, with the only changes exerted in the E-value that was adjusted to 10^−3^. Smith-Waterman optimal alignments were achieved. A Perl script was used to collect and filter the hits. To obtain non-redundant orthologs additional filters including identity and coverage of higher than 50% were applied. Using the raw sequence reads, collected genes and fragments of genes were manually extended and curated. AUGUSTUS gene prediction tool^18^ was used to identify 5’UTR and 3’UTR including introns. Nucleotide sequences were translated to protein and aligned with ortholog proteins of *Drosophila melanogaster* and *Helicoverpa armigera* in Clustal omega online tool^35^. The sequence of the receptor was checked in pfam^36^ to determine the protein family of the receptor. Accordingly, conserved residues and motifs were investigated in the sequence. The preprohormone sequence of AST-C was subjected to SignalP 5.1 version to obtain the mature peptide sequence^37^. N-terminus residues of AST-C peptides was modified (Glutamine to pyroglutamate). The modified version was synthesized by GenScript company. Peptide was dissolved in 0.1% (w/v) bovine serum albumin (BSA)-containing 1X phosphate buffered saline (PBS). 0.1% (w/v) BSA-containing PBS was used as the vehicle treatment.

### 4.2. Cell culture and transfection

HEK-TSA cells were cultured in DMEM (PAN Biotech) containing 4,5 g L^-1^ Glucose, 10% FBS, 2 mM L-Glutamine, Penicillin (50 mg mL^-1^) and Streptomycin (50 mg mL^-1^) at 37 °C in 5% CO_2_ incubator. Cells were routinely checked for mycoplasma contamination using MycoAlert™ Mycoplasma Detection Kit (Lonza). Cells were seeded in 6- or 10-cm cell culture dishes prior to transfection. Transfections were performed using Effectene Transfection Reagent (QIAGEN) according to the manufacturer’s instruction.

### 4.3. Immunocytochemistry and microscopy

cDNA of *T.pit* was used to amplify AstR-C receptor using 5’-XhoI ATGGAGCTCGAA −3’ and 5’-HindIII GAGTCGCGAATG −3’ primers. It was then cloned in pSYFP-N1 (4717bp) plasmid ^38^ to add YFP to the C-terminus of the receptor. This construct was named as AstRC-SYFP. HEK-TSA cells were seeded in 6-well plates on Poly-L-Lysine (PLL) (Sigma-Aldrich)-coated cover slips and transfected with AstRC-SYFP. Cells were washed once with pre-warmed 1X PBS 24 hours after transfection and fixed with 1 ml ice-cold 4% paraformaldehyde for 20 minutes at room temperature, and then washed 3 times with warm 1X PBS. 1X CellMask (Thermo Fisher Scientific) and Hoechst 33342 Solution (Thermo Fisher Scientific) were used according to the manufacturer’s protocols, in order to label the cell membrane and nucleus, respectively. Labeled cells on cover slips were mounted on glass slides using VectaShield Antifade Mounting Medium (Vector Laboratories). Samples were imaged using a Leica TSC SP8 confocal microscopy setup equipped with an HC PL APO 40x/1.30 Oil CS2 objective. Localization of *T.pit* AstR-C was imaged via illumination of EYFP (λex/λem: 514/518-580 nm), cell membrane was imaged via CellMask (λex/λem: 649/655-700 nm) and the nuclei were imaged via Hoechst 33342 stain (UV laser, λex/λem: 405/460-490 nm). Images were obtained with the LAS X software in a 1024 × 1024 pixel format, consisting of 4 averaged line scans. The scan speed was set to 400 Hz and pinhole was set to Airy 1.

### 4.4. G protein activation assay

cDNA of the *T.pit* AstR-C receptor was amplified using 5’-HindIII ATGGAGCTCGAAGAC-3’ and 5’-BamHI TCAGAGTCGCGAAT-3’ primers. The receptor was cloned in the mammalian expression vector, pcDNA3.1 plasmid (Invitrogen, V790-20). This construct will be referred as pc-AstR-C. Different FRET biosensors including G_i1_, G_i2_, G_i3_ ^39^, G_q_^40^, G_s_^41^ and G_13_^42^ were used to measure the G protein activation. In these biosensors G_α_ subunit is tagged with mTurquise2 and G_α2_ is tagged with mVenus^39^. Constructs were transfected transiently to HEK-TSA cells. At 50–70% confluency, cells were transfected with pc-AstR-C and FRET biosensors. Twenty-four hours later, cells were reseeded in black-bottom 96-well plates (Corning) as 75.000 per well. Twenty-four hours after reseeding, cells were subjected to FRET measurement. Before the measurement, DMEM was substituted with HBSS, and then basal FRET ratio was measured in 90 μL buffer. Subsequently, 10 μL of 10-fold ligand solution or buffer (negative control) was applied to each well and the stimulated FRET ratio was recorded. All FRET experiments were conducted at 37 °C with a Synergy Neo2 plate reader (BioTEK) equipped with 420/50 nm excitation and 485/20 nm emission filters for CFP. Acceptor emission of YFP were detected with a 540/25 nm (FRET) filter.

### 4.5. Temporal kinetics of G protein activation

The same constructs and cell culture procedure as G protein activation assay was used. During the reseeding step, cells were transferred to Poly-L-Lysine-coated coverslips in 6-well cell culture dishes. 16 hours after re-seeding, coverslips were placed in a metal chamber, washed with PBS supplemented with HBSS. Kinetics measurements were performed on a Zeiss Axiovert 200 inverted microscope equipped with an oil immersion 63x objective lens and a dual-emission photometric system. Ligand application during live FRET measurement was performed using a high-speed perfusion system (ValveLink 8.2, Automate Scientific). Cells were excited with light from a polychrome IV. Illumination was set to 40ms out of a total integration time of 100ms. Applying the excitation at 436 ± 10 nm (beam splitter DCLP 460 nm), CFP (480 ± 20 nm), YFP (535 ± 15 nm), and FRET ratio (YFP/ CFP) signals were recorded at the same time (beam splitter DCLP 505 nm). Fluorescence signals were detected by photodiodes and digitalized by an analogue-digital converter (Digidata 1440A, Axon Instruments). All data were recorded on a PC running Clampex 10.3 software (Axon Instruments). To extract the exponential time constant, tau, obtained traces were fit to a one component exponential decay function. The half-time of activation (t_1/2_) is defined as τ*ln2. In dynamic experiments, cells were stimulated with *T.pit* AST-C ligand.

### 4.6. Kinetics of receptor/G protein interaction (G-protein recruitment)

AstRC-SYFP, G_i2_ biosensor in which G_α_ was tagged with CFP were used for transient expression of the AstRC-SYFP and the G protein subunits. 1.5 × 10^6^ HEK-TSA cells were seeded onto a 55 mm dish and transfected 24 hours later. Kinetics measurements were performed as explained in “Temporal kinetics of G protein activation”.

### 4.7. βarrestin recruitment assay

AstRC-SYFP, GRK_2_ (G protein receptor kinase 2) and βarrestin2-Nluc^43^ were transfected to HEK-TSA cells. Cells were washed to substitute DMEM with the experimental buffer and incubated with the substrate of Nluc, furimazine (1:1000 of 90 μL HBSS) for 2–5 min at 37 °C. Following the incubation step, the basal BRET ratio was measured. Then, 10 μL of 10-fold ligand solution or buffer was applied to each well and the stimulated BRET ratio was recorded for 20 minutes. BRET experiments were performed at 37 °C with Synergy Neo2 (BioTEK) plate reader equipped with a 460/40 nm filter to select the NanoLuc emission.

### 4.8. Homology modeling

SWISS-MODEL online tool (https://swissmodel.expasy.org)^44^ was used to build tertiary structure of AstR-C. Different templates (PDB ID: 4N6H, 5C1M and 6DDE) with high sequence identity and similarity to AstR-C of *T.pit* were evaluated, and different models were built. The constructed models were primarily evaluated using the QMEAN ^44^ and Ramachandran plot. The acceptable models were then applied to 25-ns MD simulations, and RMSD and RMSF changes were monitored during the MD simulation time. N-terminus was built using I-TASSER webserver (https://zhanglab.ccmb.med.umich.edu/I-TASSER) ^45^ and added to the finally selected model. The ligand structure (AST-C) was also built using I-TASSER. The distance between the two Sulfur atoms of Cysteine residues were set to be kept at 2.05 Å in order to have the disulfide bond between these two residues. Five models were generated and the best one according to the C-score was chosen. C-score is at the range of [-5,2]. Bigger numbers show higher-quality models.

### 4.9. Protein Preparation

“Protein Preparation” module of the Maestro molecular modeling package (Schrödinger Suite 2017 Protein Preparation Wizard; Schrödinger, LLC, New York, 2017; Impact, Schrödinger, LLC, New York, 2017) was used to prepare both the receptor and ligand prior the MD simulations^46^. The protein refinement and minimization were performed in this step. Prime^47^ module of Maestro (Prime, Schrödinger, LLC, New York, NY, 2017) was used to resolve any problem regarding the protein structure such as missing hydrogen atoms, side chains, loops or flipped residues. The protonation states at pH 7.4 was assigned using PROPKA^48^. OPLS2005 force field^49^ was used for the minimization and optimization processes.

### 4.10. System Preparation

G_α_ part of Gi complex available at protein data bank (PDB) (PDB ID, 6DDE) was taken and aligned at the intracellular part of the receptor to be merged. To fill the gaps available in the resolved structure of G_α_, “Crosslink Proteins” module of Schrödinger2015 (Schrödinger Release 2015-2: Prime, S., LLC, New York, NY, 2015) was used. Simple *de novo* loop creation was chosen for the linker conformation prediction, and implicit solvent energy calculation of Prime module was selected for the energy calculation. The orientation of the constructed models for AstR-C in the membrane was adjusted using the Orientations of Proteins in Membrane (OPM) database^50^. The “Desmond System Builder” module of Maestro was used to set up the biological system which consists of solvent, membrane, counter ions and water. The protein was embedded in POPC (1-palmitoyl-2-oleoyl-sn-glycero-3-phosphocholine) lipid bilayer and TIP3P explicit water ^51^ was selected. 0.15 M NaCl was added to the system.

### 4.11. Molecular Dynamics (MD) Simulations

Desmond package was used for the MD simulations (Desmond Molecular Dynamics System, D. E. Shaw Research, New York, NY, 2017). OPLS 2005 force field^49^ was used. The equilibration step was performed using the default algorithm of Desmond. The simulations were run at 300 K, which is the recommended temperature when using POPC lipid bilayer. To keep the temperature at 300 K and the pressure at 1.01325 bar, Nose-Hoover thermostat^52^ and Martyna-Tobias-Klein method ^53^ were applied to the system. The particle mesh Ewald method^54^ was applied to calculate the long-range electrostatic interactions. For both van der Waals and Coulombic interactions, a cut-off radius of 9.0 Å was used. The time-step was assigned as 2.0 fs. NP_γ_T ensemble was used during the production step of MD simulations with surface tension of 4000 bar/Å as it is the recommended surface tension for NP_γ_T ensemble. AstR-C/ST-C system was subjected to 500 ns MD simulation and three independent replica simulations were performed. The trajectory files collected during MD simulations were used for the analysis. RMSD and RMSF of the complexes were analyzed during the MD simulation time. 100 trajectory frames were recorded and MM-GBSA binding free energies of AST-C to AstR-C was calculated. VSGB 2.0 solvation model at Prime module of Maestro was utilized during MM/GBSA calculations.

### 4.12. Molecular Docking Studies

ClusPro web server^55^ (http://nrc.bu.edu/cluster) was used for the docking studies. A mask file including the repulsion site was provided to the program. The contributing residues of the receptor in the ligand-protein interaction were evaluated by in “Ligand Interaction Diagram” of the Maestro package.

### 4.13. *In silico* Binding Pocket Verification

A representative structure with minimum RMSD value was selected from trajectories of 500-ns MD simulations done for apo AstR-C. The desired residues were substituted with Ala and mutant receptors were subjected to 200-ns MD simulations. Representative structure with the minimum RMSD value was chosen at each system and docking with the native ligand was applied. The best docking pose was selected, and two different systems were generated for 200-ns MD simulations. In the first one, the obtained pose was directly used in MD simulations. In the second approach, however, G_α_ subunit was inserted in the intracellular interface of the receptor and then system was prepared for MD simulations. The effect of the mutations on the state (active, intermediate, and inactive) of the receptor was investigated measuring the *Δd* that is calculated as given in Equations 1-3.

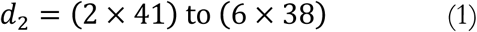

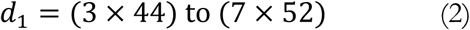

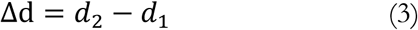

d_1_ and d_2_ obtain by measuring the distance between specific residues. In the given formula, residues are specified by generic numbering offered by gpcrdb. In AstR-C, d_2_ is the distance between M75^2×41^ and L254^6×38^ and d_1_ is the distance between C129^3×49^ and L308^7×52^. *Δd* was measured along the MD simulation time using the “Simulation Event Analysis” module of Schrodinger.

### 4.14. *In vitro* Binding Pocket Verification

Q5® Site-Directed Mutagenesis Kit (NEB, Beverly, MA) was used to substitute the selected residues to Ala in the pc-AstR-C construct. G protein activation assay was performed to evaluate the effect of each substitution on the receptor activation.

### 4.15. Principal Component Analysis (PCA)

Large-amplitude motions along MD simulations could be extracted using a dimensional reduction method called PCA. The details of algorithm and how to apply this method for MD simulations could be found in other papers^56, 57, 58^. In this study Bio3D package of Grant *et al*.^59^ was utilized using R program. MD trajectories obtained from different independent replica simulations were concatenated to increase the number of conformations for protein. All the frames of trajectories were aligned with respect to an initial reference state before performing PCA to eliminate translational and rotational motions of protein and just to focus on internal fluctuations. Only alpha-C (-) atoms of proteins were used for PCA to focus on backbone movements. Here, we have applied PCA for both AstR-C and G_α_ protein separately. We have performed PCA for both WT and Q271A mutated systems, for which MD simulations was extended to 500-ns, to elucidate the effect of mutation in addition to determine the overall combined motions of proteins. Both holo and apo forms of systems were considered to elucidate the effect of ligand binding on receptor.

### 4.16. Dynamics Cross Correlation Analysis

The cross-correlation between atomic fluctuations/displacements are useful to provide information about the effect of mutations, ligand-binding etc. on the receptor/protein structure^58^. Here, Bio3D package in R program was used and C_α_ atoms of proteins were utilized to focus on backbone of proteins. For both receptor AstR-C and G_α_ proteins cross correlation analysis were performed in four different systems for which PCA also applied. Dynamic cross-correlation maps (DCCM) were plotted to visualize the correlation between residues in Bio3D package.

### 4.17. Data Analysis and Visualization

ImageJ (National Institute of Health) was used to analyze the raw microscopy images. Further processing of the data was done in Excel (Microsoft Office). All concentration-response data were fitted using nonlinear regression models with Prism 6 (GraphPad Software, San Diego, CA, USA). For each concentration, the response is normalized to the buffer only dataset. Visual Molecular Dynamics (VMD) software^60^ was used for visualization and image generation.

## Accession codes

The coding sequence of AstR-C and AST-C were deposited on GenBank under MN871948 and MT254058 accession number, respectively.

## Supporting Information Available

This material is available free of charge via the Internet.

## Notes

The authors declare no competing financial interest.

## Abbreviations

AstR-C: allatostatin receptor type C
GPCRs: G protein-coupled receptors
AST-C: allatostatin C
JH: Juvenile hormone
*T.pit*: *Thaumetopoea pityocampa*
WGS: whole genome sequencing
RET: Resonance energy transfer
3D: Three-dimensional
MD: molecular dynamics
ECL: extracellular loop
ICL: intracellular loop
FRET: Förster resonance energy transfer
BRET: Bioluminescence resonance energy transfer
YFP: yellow fluorescent protein
CFP: cyan fluorescent protein
C_α_: Carbon alpha
RMSD: root mean square deviations
RMSF: root mean square fluctuations
Ala: Alanine
DCCM: dynamical cross-correlation map
PDB: protein data bank
BSA: bovine serum albumin
PBS: phosphate buffered saline
MM/GBSA: Molecular Mechanics/ Generalized Born Surface Area.

## Acknowledgment

This study is supported by funding from TUBITAK-119Z921 (The Scientific and Technological Research Council of Turkey) and COST ACTION ERNEST (Ernest CA18133).

---

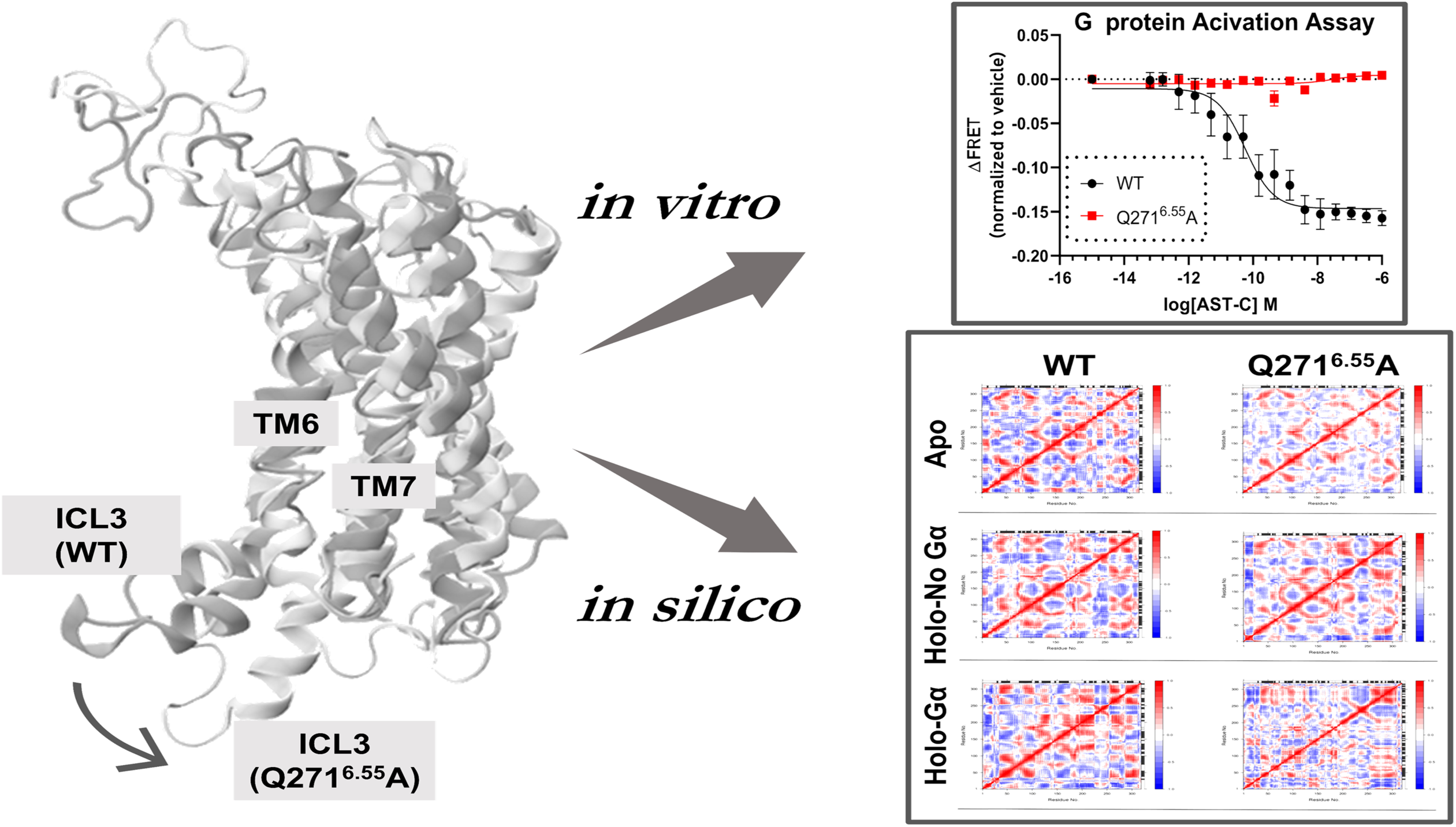

