## Supplemental Files for "Structural and functional characterization of allatostatin receptor type-C of *Thaumetopoea pityocampa* revealed the importance of Q271^6.55^ residue in G protein-dependent activation pathway"

**Supporting Information**

**A**

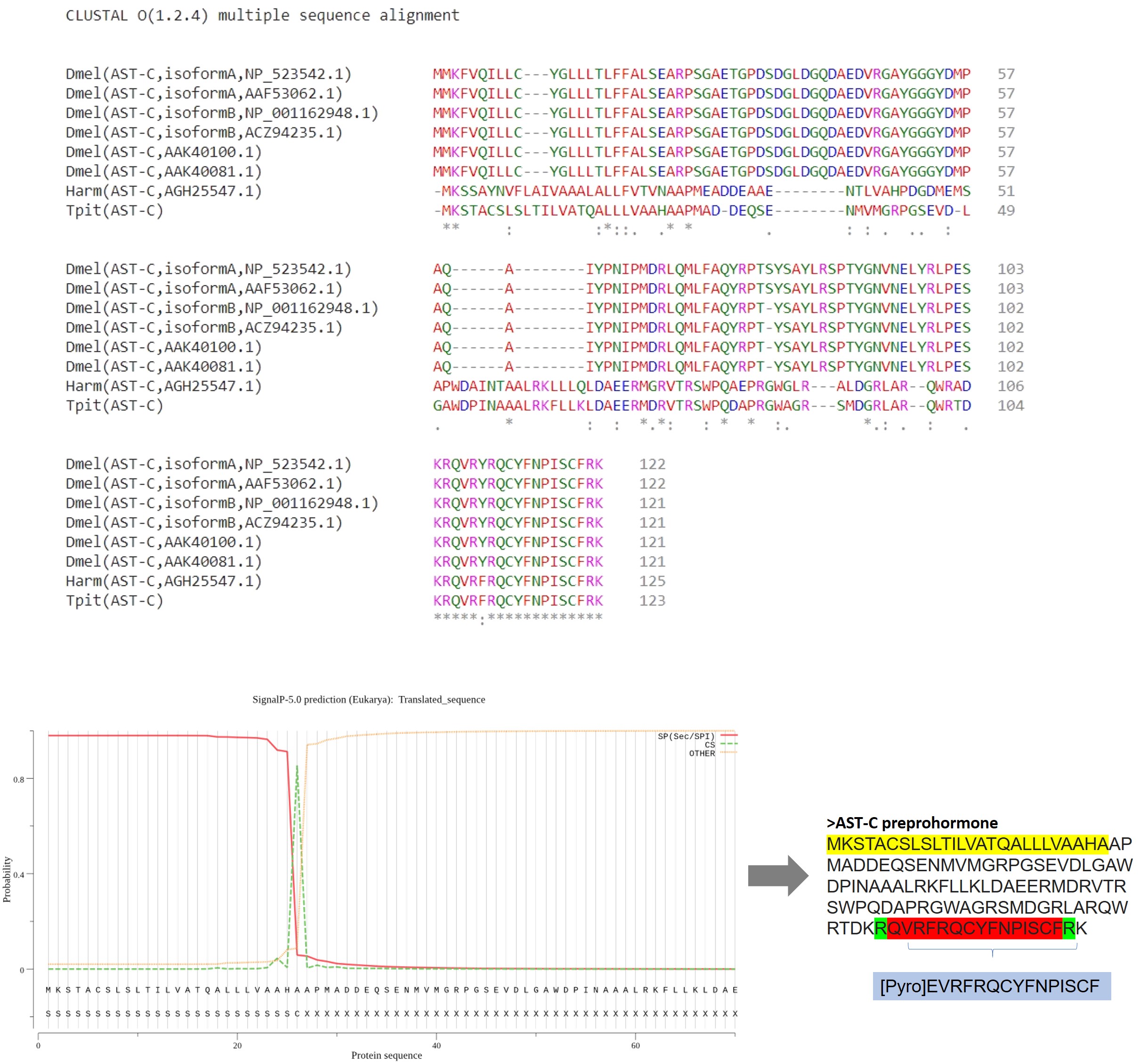

**B**

**Supplementary Figure S1.** AST-C preprohormone. (A) Alignment of the AST-C preprohormone sequence of *Drosophila melanogaster* (shown as Dmel) and *Helicoverpa armigera* (shown as Harm) in clustal omega (<https://www.ebi.ac.uk/Tools/msa/clustalo/>). (B) SignalP-5.0 was used to predict the signal peptide. In the left side, Probability vs. Protein sequence plot is given, in which the red line represents a part of the sequence which comprises the signal peptide (SP), green line shows the cleavage site (CS) and yellow line denoted as “OTHER” depicts the probability that the sequence does not have any kind of signal peptide. Result showed that there is a cleavage site (CS) between the 26^th^ and 27^th^ residues, AHA-AP, with the probability of 0.8541. The signal peptide is highlighted in yellow. The proteolytic dibasic cleavage, highlighted in green, was applied according to the rules suggested by Veenstra, 2000. The resultant 15-amino acid peptide highlighted in red further modified in the N-terminus. Glutamine is converted to pyroglutamate. The mature peptide sequence is highlighted in blue.

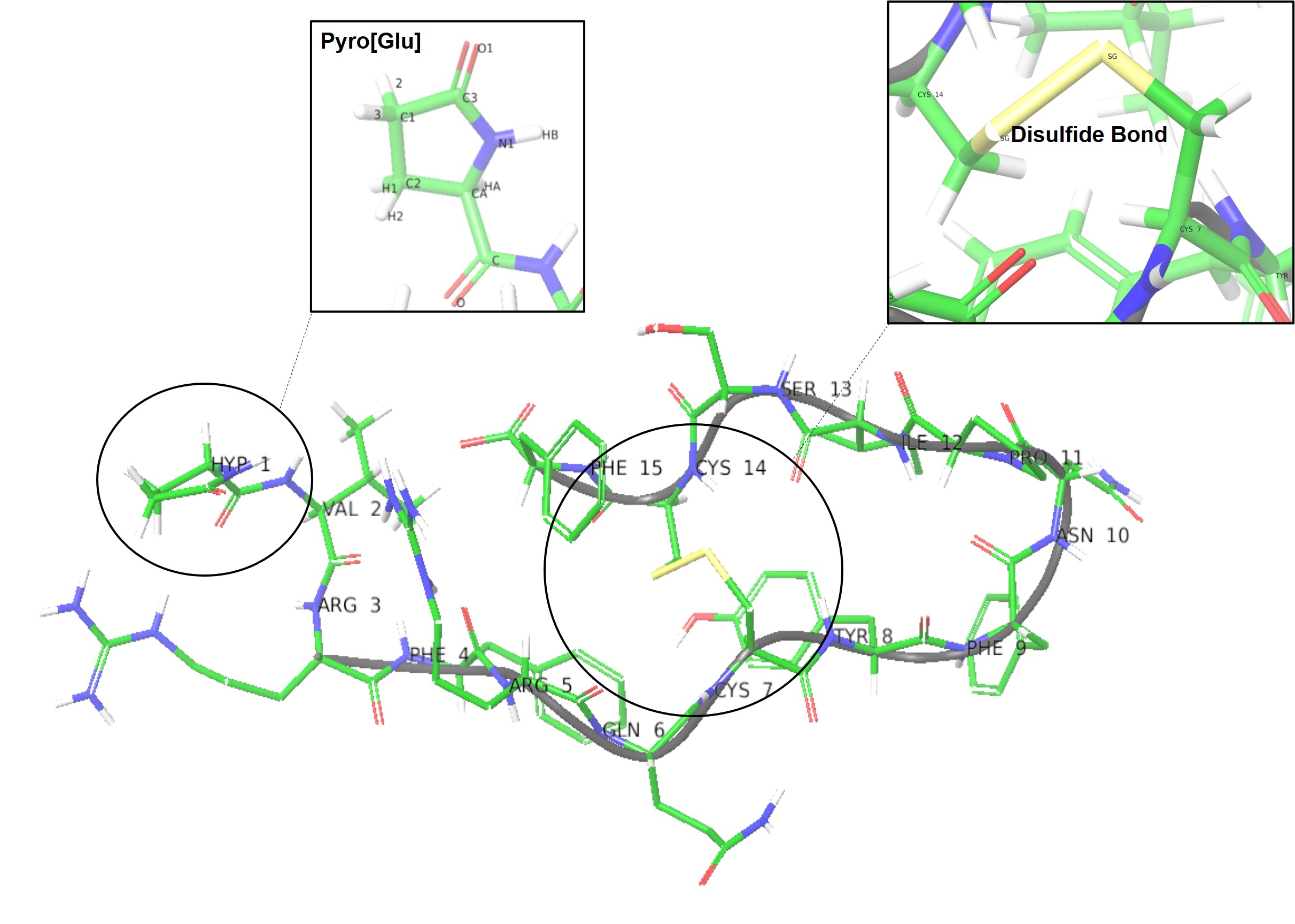

**Supplementary Figure S2**. AST-C structure. The first residue of the ligand was modified to (Pyro[Glu], and disulfide bond was formed between the Sulfur atoms of Cys7 and Cys14.

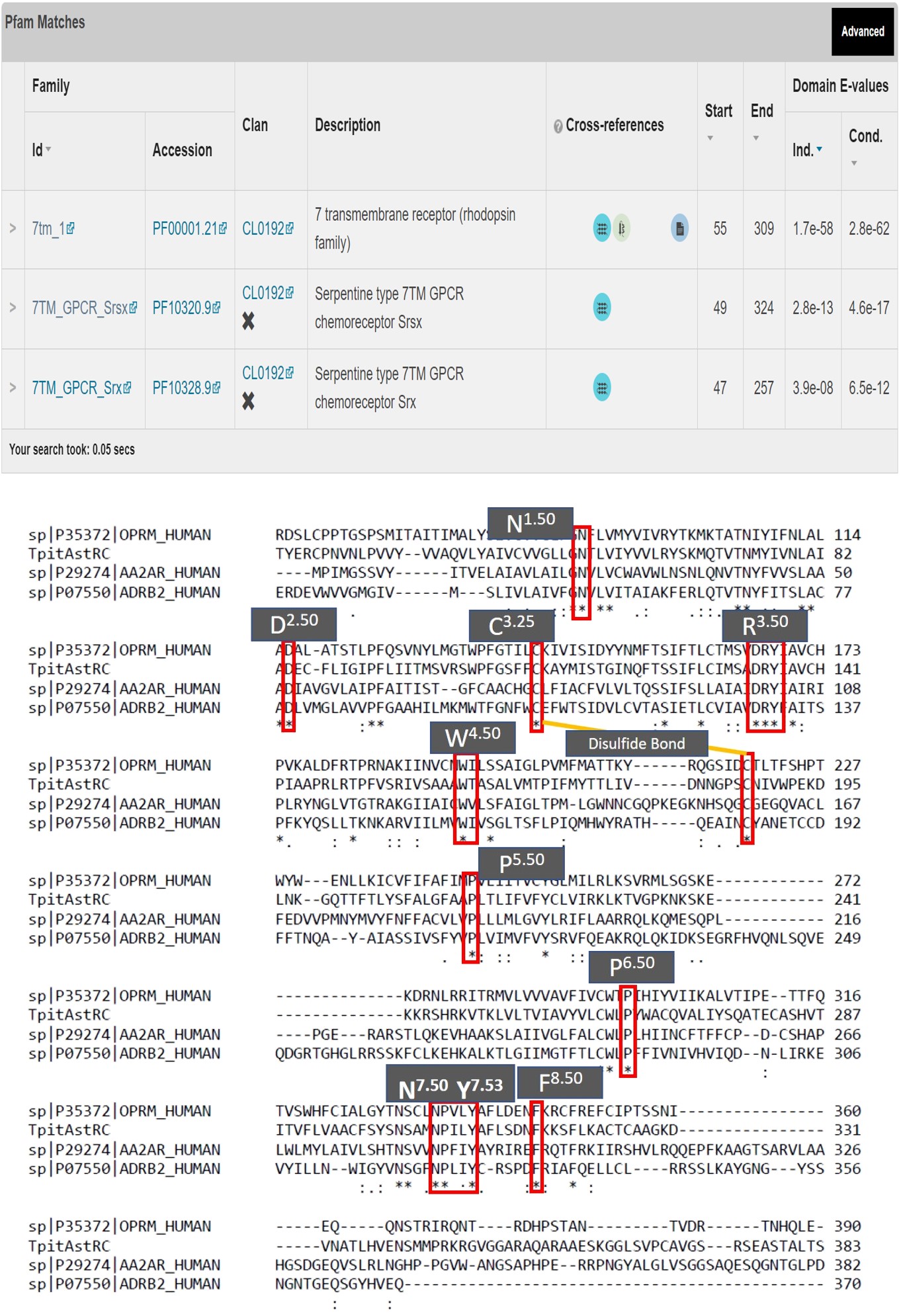

**B**

**A**

**Supplementary Figure S3.** Sequence analysis of *T.pit* AstR-C. A) Analysis in pfam using Hmmscan. The receptor is a 7 transmembrane receptor belonging rhodopsin family GPCRs (<https://pfam.xfam.org/>). B) Alignment of the protein sequence of well-known Class A GPCRs, including Beta-2 adrenergic receptor (UniprotKB: P07550), mu opioid receptor (UniprotKB: P35372) and adenosine A2a receptor (UniprotKB: P29274) in clustal omega (<https://www.ebi.ac.uk/Tools/msa/clustalo/>).

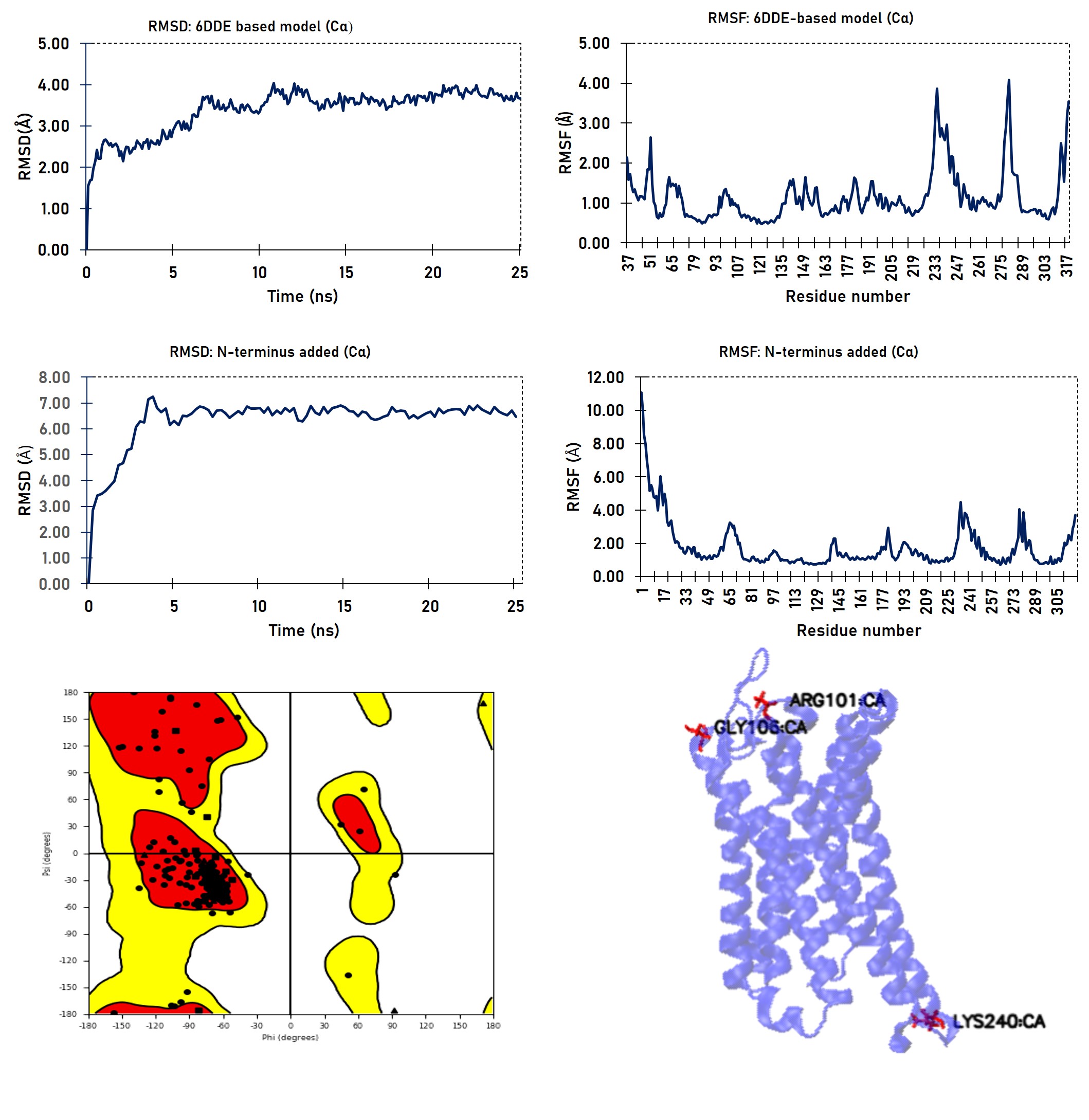

**C**

**A**

**F**

**E**

**B**

**D**

**Supplementary Figure S4.** Model analysis. The quality of the constructed model was evaluated considering RMSD and RMSF changes and Ramachandran plot. A) RMSD changes during 25 ns MD simulation time. B) RMSF changes of the residues of AstR-C during 25 ns MD simulation time. C) RMSD changes of the model after addition of the N-terminus during 25 ns MD simulation time. D) RMSF changes of the residues of the model after addition of the N-terminus during 25 ns MD simulation time. E) Ramachandran plot of the model (Red: favored region, Yellow: allowed region). F) Receptor localization of outlier residues, Arg101, Gly106 and Lys204.

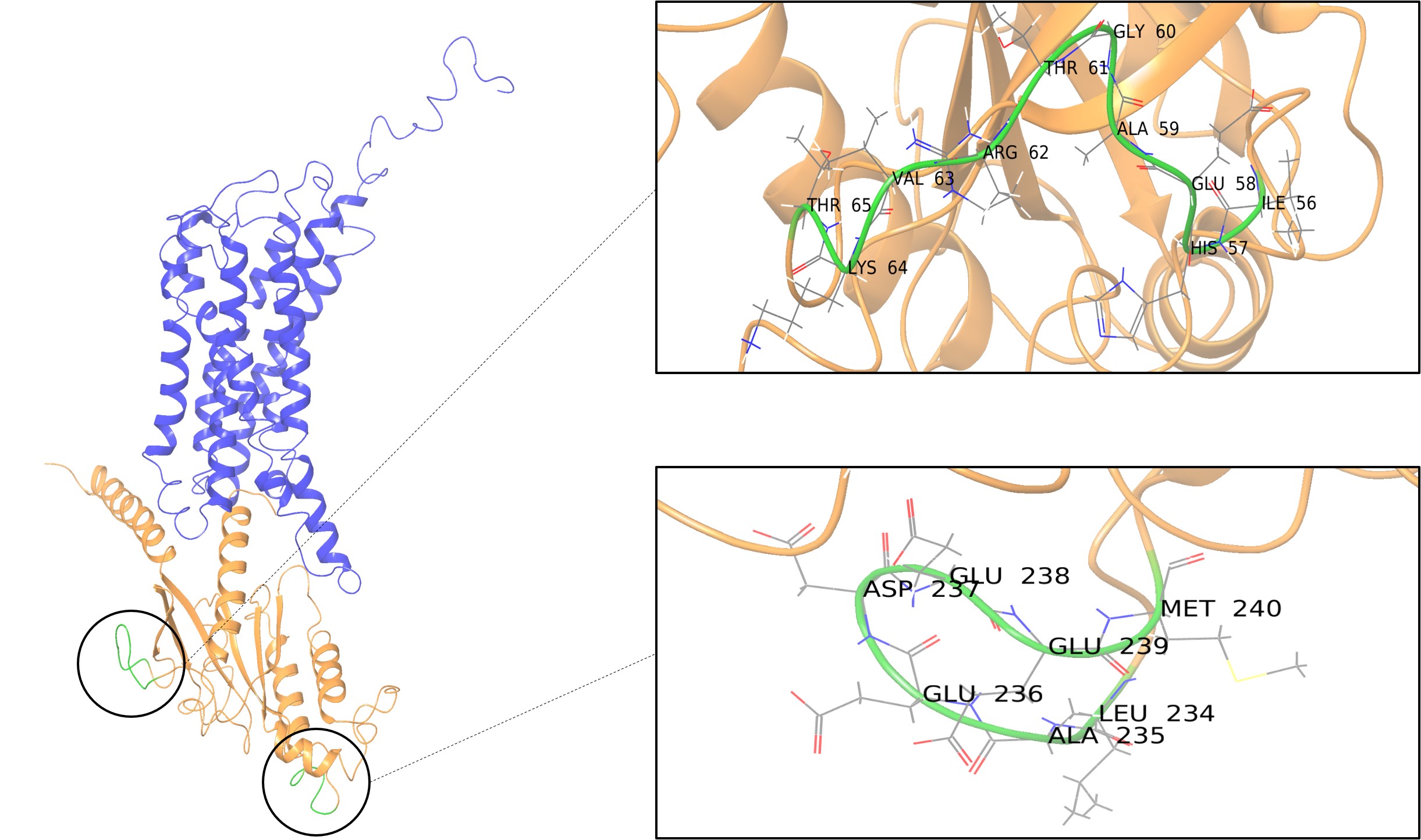

**Supplementary Figure S5.** Crosslinking the gaps. Gaps in G_α_ part of the structure with PDB ID, 6DDE were filled and crosslinked.

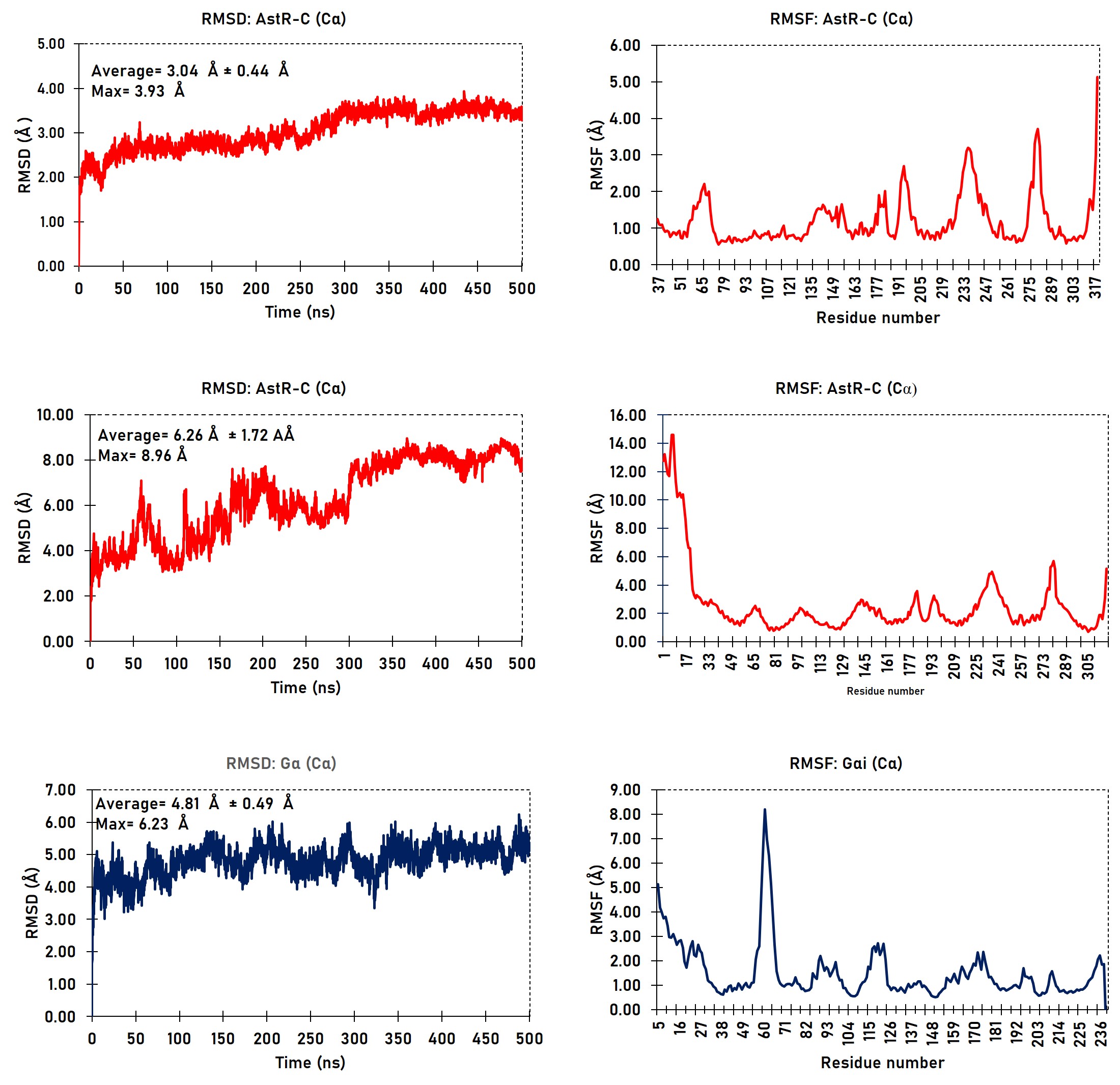

**F**

**D**

**C**

**E**

**B**

**A**

**Supplementary Figure S6.** System stability during the simulation time. (A) RMSD and (B) RMSF changes of AstR-C when N-terminus is not considered in calculations. (C) RMSD and (D) RMSF changes of AstR-C when N-terminus is considered. (E) RMSD and (F) RMSF changes of Gα. MD simulation time was 500 ns. The analysis is given for Carbon-alpha (C_α_).

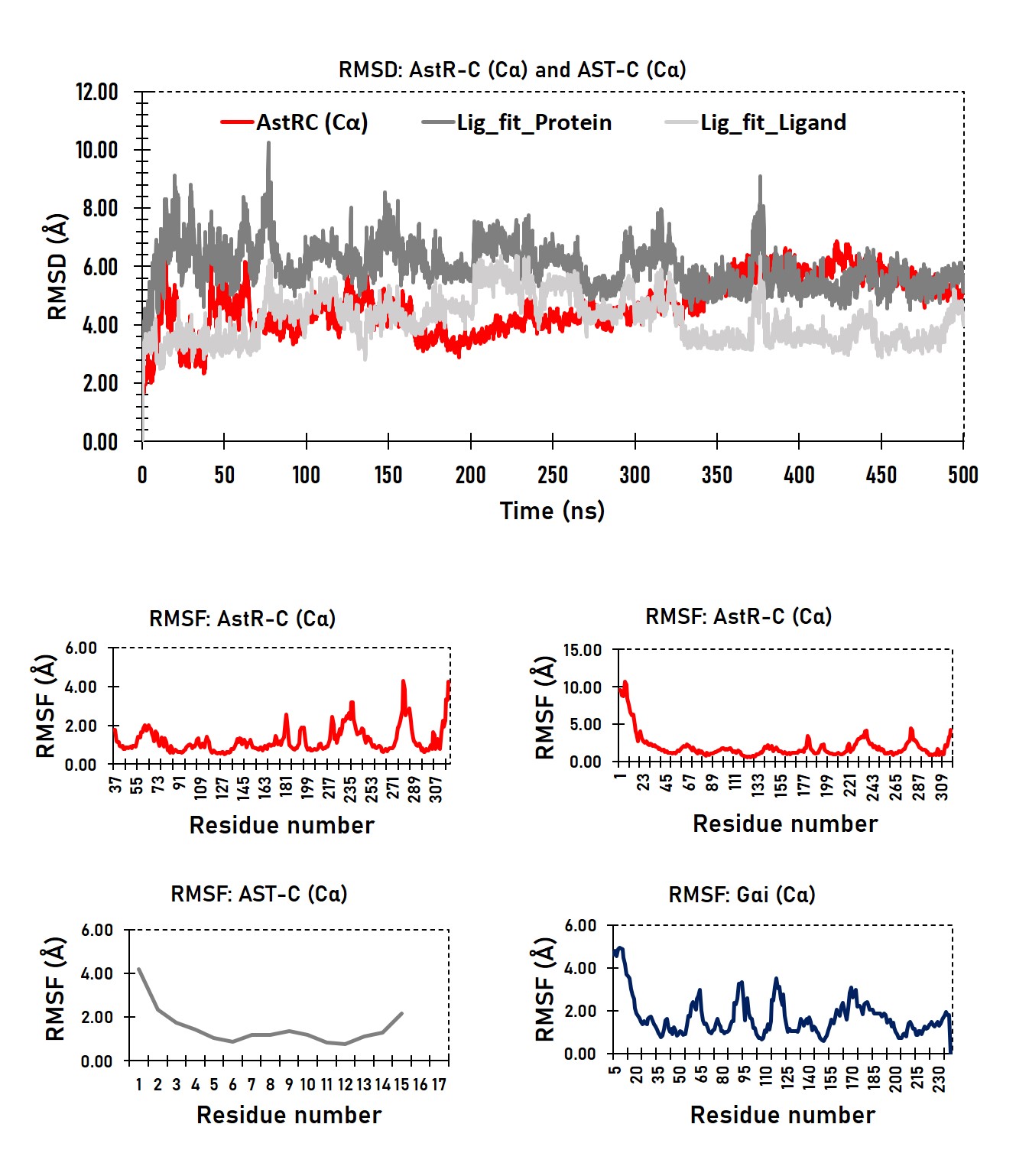

**D**

**A**

**B**

**E**

**C**

**Supplementary Figure S7.** Stability of the best Protein-Ligand pose during 500 ns MD simulation time. (A) RMSD changes of AstR-C (shown in red) and AST-C when fitted to the receptor’s first frame denoted as “Lig-fit-Protein” (shown in dark gray) and AST-C fitted to the ligand’s first frame, denoted as “Lig-fit-Ligand” (shown in light gray). RMSF changes of (B) AstR-C without N-terminus, (C) with N-terminus, (D) AST-C. (E) G_αi_. (Note: The analysis is given for Carbon-alpha (Cα)).

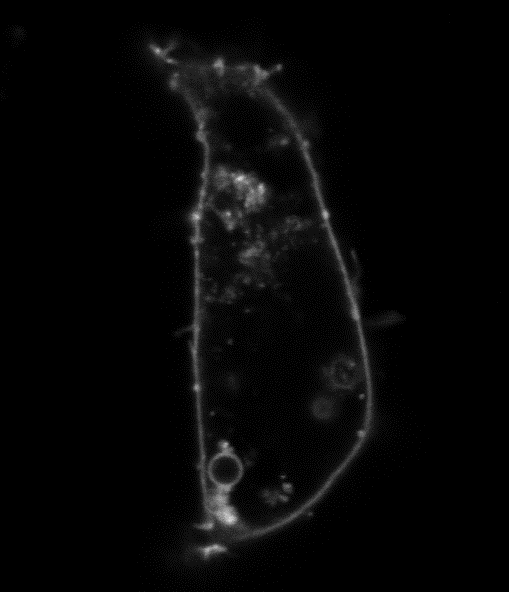

**Supplementary Figure S8.** Cell localization of *T.pit* AstR-C with Ala substitution at Q271^6.55^. Receptor with a Point mutation of Q271A is majorly expressed in cell membrane.

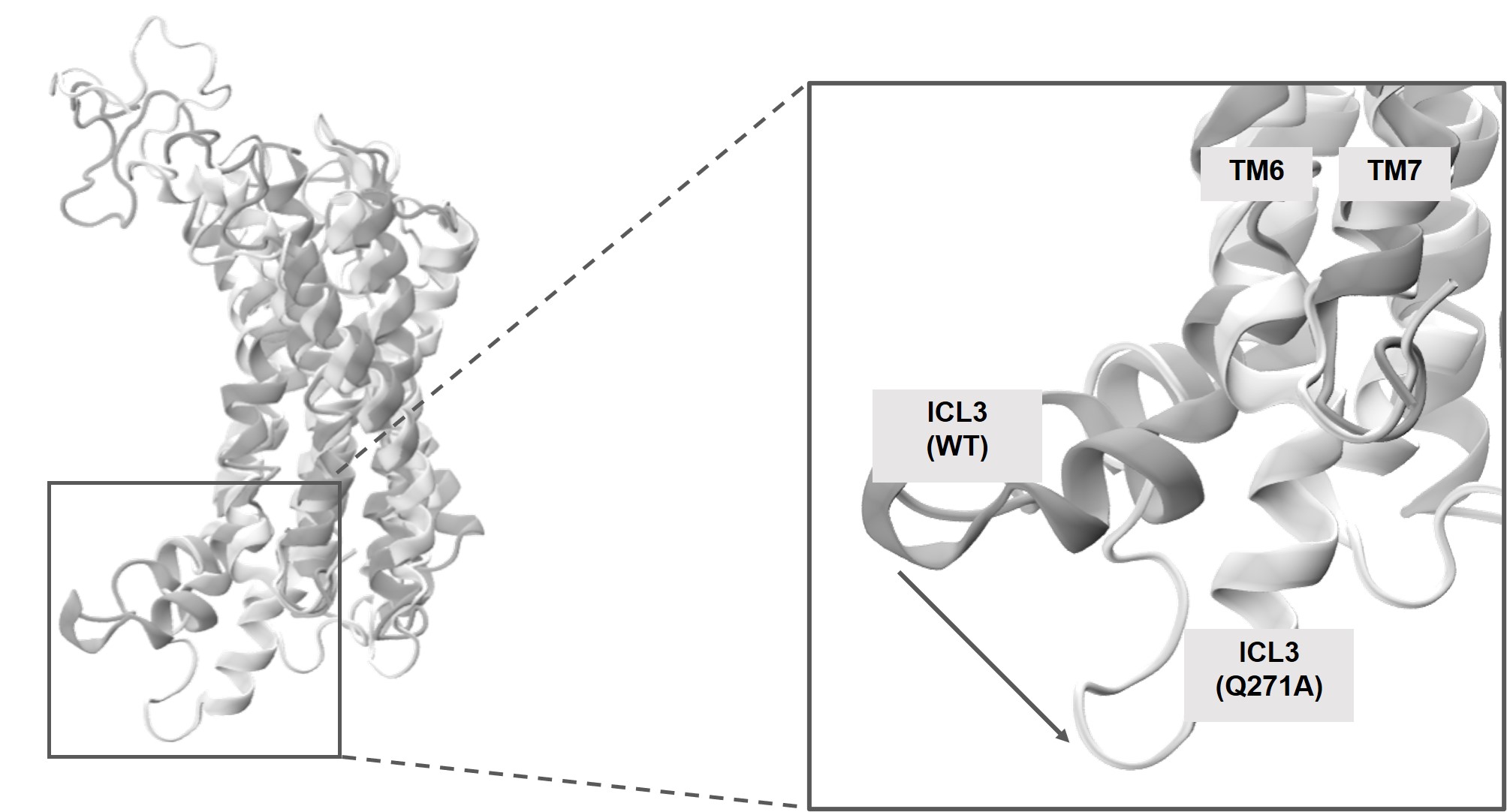

**Supplementary Figure S9.** Superimposition of WT and Q271A mutant receptor at Apo form. A distict positioning of ICL3 was observed for Q271A mutant (light gray) when compared to WT (dark gray).

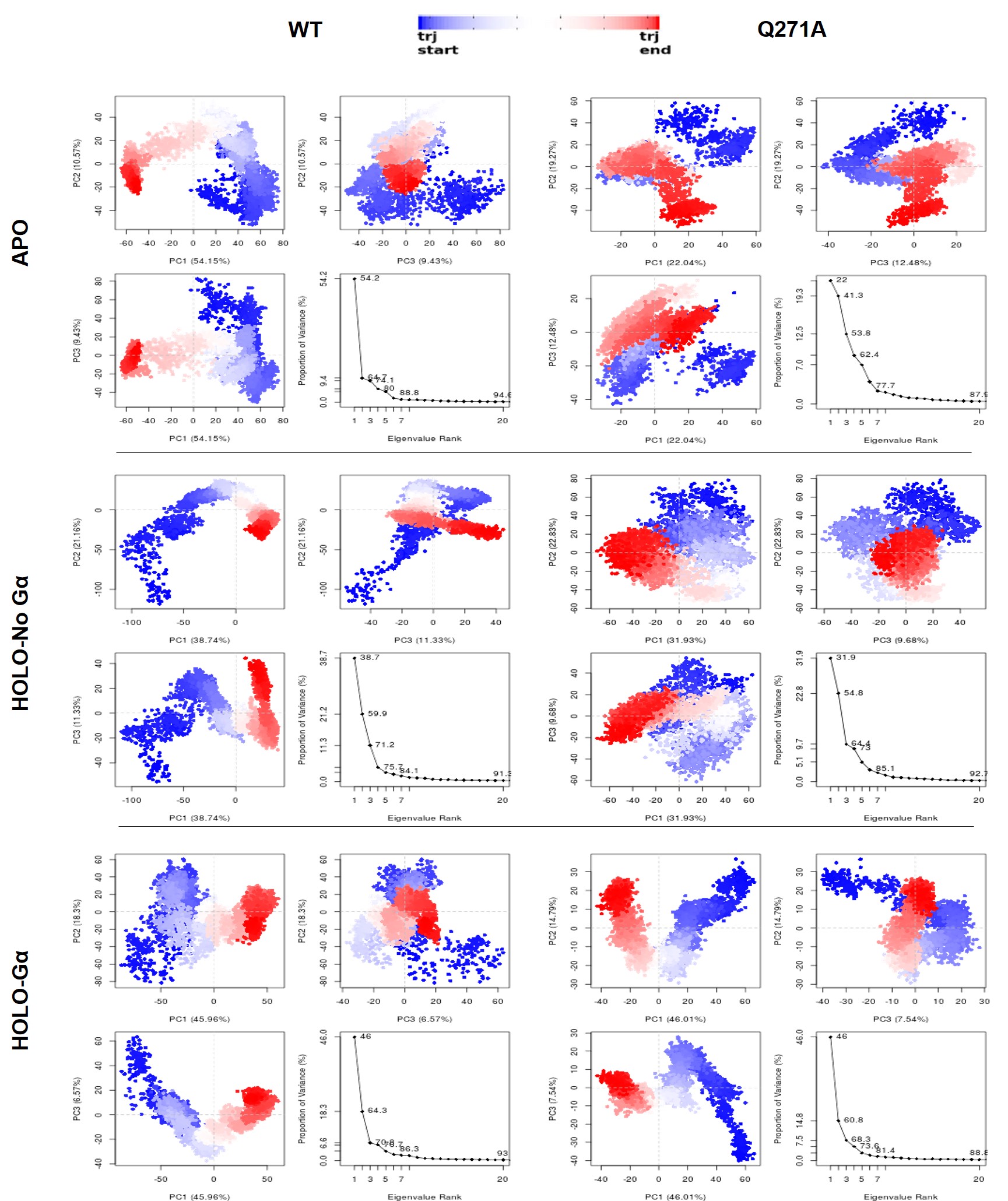

**Supplementary Figure S10.** PCA results for trajectories with instantaneous conformations. The total mean square displacement (or variance) of atom positional fluctuations were projected onto the subspace defined by the largest three principal components. Scree plots (degree of explained variance) for all systems were also displayed to show that three dimensions were enough to capture at least 60% of the atomic motions.

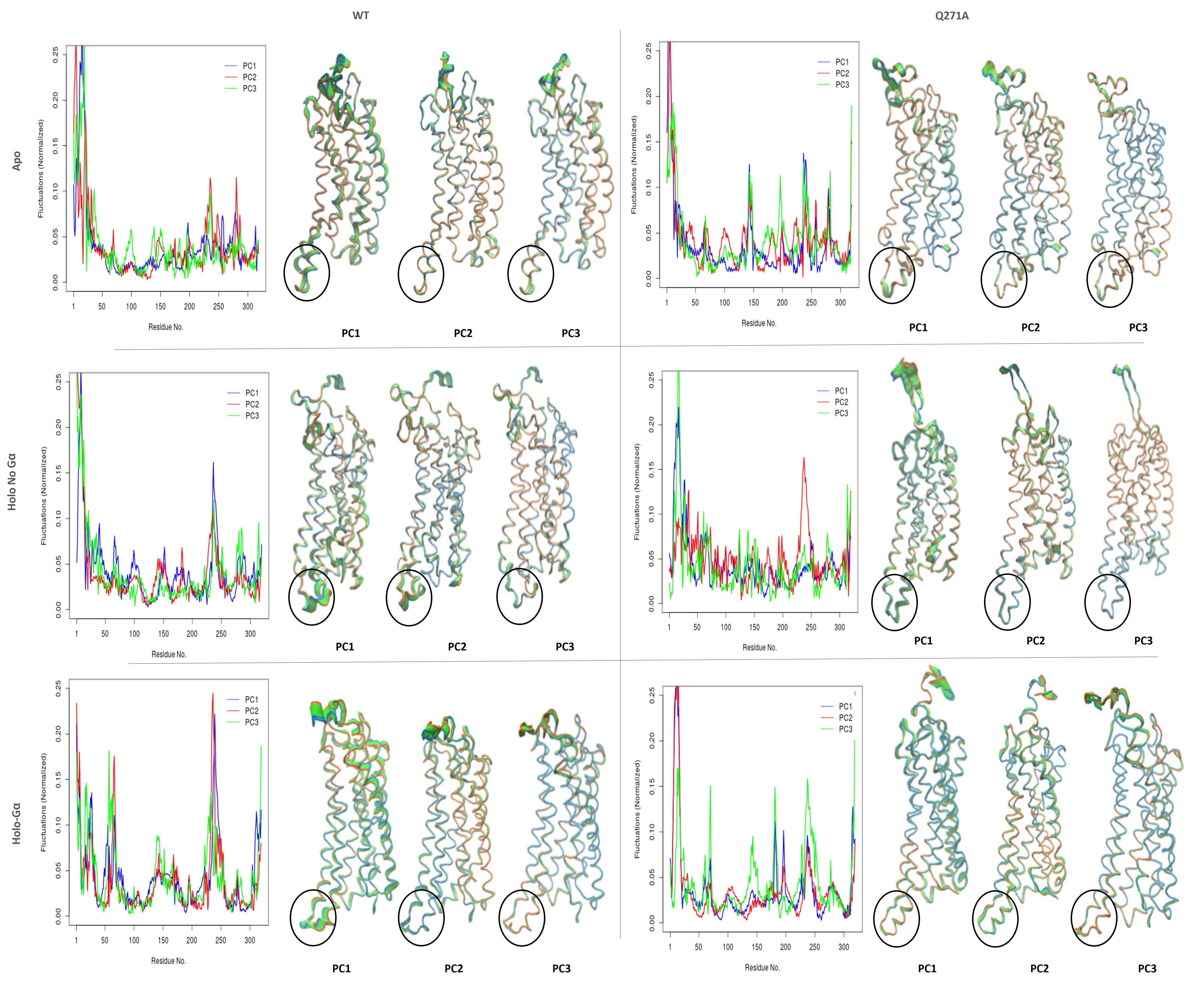

**Supplementary Figure S11**. Representation of first three PCs for WT and Q271A at Apo and Holo with/without Gα. Blue-Green-Red colors indicate order or increasing mobility (effects of amino-acids on the specific PC)**.** Broadening tubes also indicate higher mobility as they are based on trajectory frames**.** ICL3 is denoted by circles in each PC.
